# Levelling out differences in aerobic glycolysis neutralizes the competitive advantage of oncogenic *PIK3CA* mutant progenitors in the esophagus

**DOI:** 10.1101/2021.05.28.446104

**Authors:** Albert Herms, Bartomeu Colom, Gabriel Piedrafita, Kasumi Murai, Swee Hoe Ong, David Fernandez-Antoran, Christopher Bryant, Christian Frezza, Bart Vanhaesebroeck, Philip H. Jones

**Author notes:** These authors contributed equally to this work (Albert Herms, Bartomeu Colom).

## Abstract

Normal human tissues progressively accumulate cells carrying mutations. Activating mutations in *PIK3CA* generate large clones in the aging human esophagus, but the underlying cellular mechanisms are unclear. Here, we tracked mutant *PIK3CA* esophageal progenitor cells in transgenic mice by lineage tracing. Expression of an activating heterozygous *Pik3ca^H1047R^* mutation in single progenitor cells tilts cell fate towards proliferation, generating mutant clones that outcompete their wild type neighbors. The mutation leads to increased aerobic glycolysis through the activation of Hif1α transcriptional targets compared with wild type cells. We found that interventions that level out the difference in activation of the PI3K/HIF1α/aerobic glycolysis axis between wild type and mutant cells attenuate the competitive advantage of *Pik3ca^H1047R^* mutant cells *in vitro* and *in vivo*. Our results suggest that clinically feasible interventions that even out signaling imbalances between wild type and mutant cells may limit the expansion of oncogenic mutants in normal epithelia.

## Introduction

As humans age, normal tissues accumulate clones carrying somatic mutations (Blokzijl et al., 2016; Lee-Six et al., 2018; Martincorena et al., 2018 ; Martincorena et al., 2015; Moore et al., 2020; Yizhak et al., 2019; Yokoyama et al., 2019; Yoshida et al., 2020). An example is the esophagus, in which the normal epithelium develops into a patchwork of mutant clones by middle age (Martincorena et al., 2018). There is genetic evidence that the most prevalent mutations are in genes under strong positive selection, arguing that they confer a proliferative or survival advantage over wild type cells (Hall et al., 2019). However, little is known about the cellular mechanisms that underpin the clonal expansions triggered by most of the mutant genes. Understanding these processes is important as it both reveals genes that regulate progenitor cell behavior and, as most of the selected mutant genes are frequently mutated in cancer, gives insight into the earliest stages of neoplastic transformation. The identification of pharmacologically-druggable mutations may also open opportunities for cancer prevention.

One gene that is recurrently mutated in normal human esophagus is *PIK3CA,* which encodes the p110α catalytic subunit of phosphoinositide 3-kinase (PI3K) (Martincorena et al., 2018; Yizhak et al., 2019; Yokoyama et al., 2019). PI3K is a signaling hub activated by insulin and growth factors to regulate a broad range of processes including cell proliferation, survival, growth and metabolism, mainly through the activation of the Akt/mTOR signaling axis (Fruman et al., 2017; Madsen et al., 2018). Activating *PIK3CA* mutations are recurrently found in solid tumors including esophageal squamous cell carcinoma (ESCC), and in overgrowth syndromes and vascular malformations (Castel et al., 2016; Castillo et al., 2016; Consortium, 2017; Madsen et al., 2018).

Reviewing published data, 74 missense *PIK3CA* mutant clones were identified in 18 cm^2^ of histologically normal human esophageal epithelium (Martincorena et al., 2018). *PIK3CA* missense mutations generate particularly large clones in the human esophagus (**Figure 1A**). Indeed, *PIK3CA* is the mutant gene with the second highest average Variant Allele Fraction (VAF) (**Figure 1B**). The VAF corresponds to the proportion of sequence reads detected for each DNA variant and, in diploid cells, is proportional to the size of the mutant clones carrying the mutation in the sample analyzed. These mutant clones were significantly enriched for pathogenic or likely pathogenic *PIK3CA* activating mutations according to the ClinVar mutation database (https://clinvarminer.genetics.utah.edu/variants-by-gene/PIK3CA); with 47% (35/74) activating *PIK3CA* mutations observed, well above the 1% expected under neutrality (p=2e^−27^, two tailed binomial test, see methods) (Martincorena et al., 2018). Notably, mutations classified as pathogenic or likely pathogenic had significantly higher clone size or VAF than the remaining *PIK3CA* missense mutants or the total 603 silent mutations in all tested genes, which have no effect on cell behavior (**Figure 1C**). The most prevalent pathogenic mutation was *PIK3CA^H1047R^* (35% (8/23) of all pathogenic mutation events) (**Figure 1D**), which had a significantly higher VAF than the other *PIK3CA* missense mutations (**Figure 1E**). *PIK3CA^H1047R^* mutation occurs recurrently in human esophageal and other cancers (https://cancer.sanger.ac.uk/cosmic/gene/analysis?ln=PIK3CA), benign overgrowth syndromes and in vascular malformations (Cancer Genome Atlas Research et al., 2017; Castillo et al., 2016; Fruman et al., 2017; Keppler-Noreuil et al., 2014; Madsen et al., 2018). Taken together, these findings indicate that activating *PIK3CA* mutants, and particularly *PIK3CA^H1047R^*, may drive large clonal expansions in normal human esophagus.

**Figure 1.**
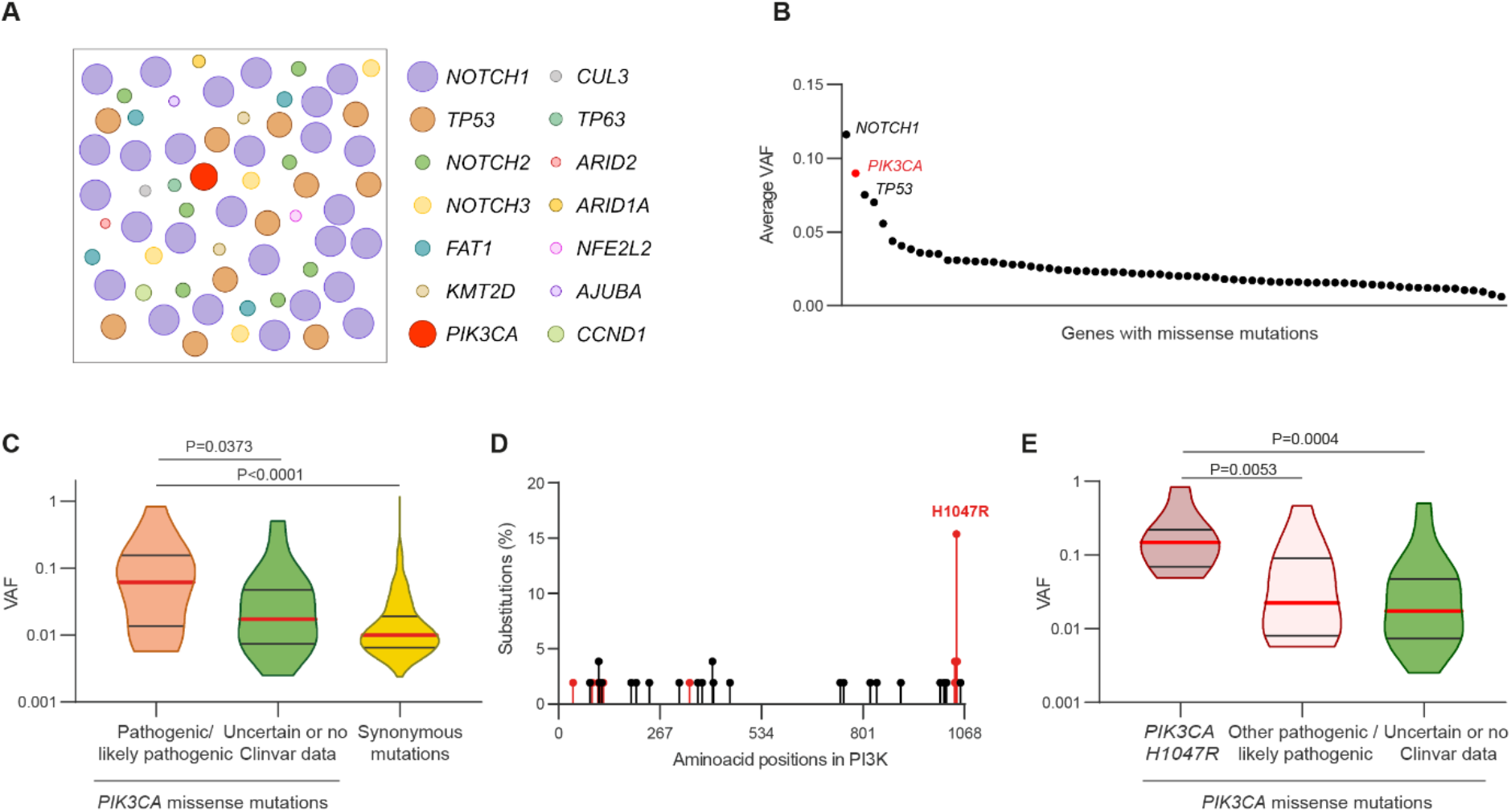
*PIK3CA^H1047R^* clones have high variant allele frequency in the human esophageal epithelium. Mutant clones in normal human esophagus (data from (Martincorena et al., 2018)).(A) Schematic representation of clones carrying missense mutations under positive selection. Clones are represented as circles, with their frequency and size relative to the mutation frequency and their variant allele frequency (VAF), respectively. VAF is proportional to clone size in diploid normal tissue. (B) Missense mutations detected more than once ordered by decreasing average VAF. VAF of missense mutations in the *PIK3CA* gene are highlighted in red. (C) Distribution of *PIK3CA* missense mutations classified according to the Clinvar database (https://www.ncbi.nlm.nih.gov/clinvar/) into pathogenic/likely pathogenic or uncertain significance/no data available. VAF distribution of the synonymous passenger mutations in all genes is also shown. Medians (red) and quartiles (grey lines) are represented. Two-tailed Mann Whitney test. (D) Distribution of codons altered by missense mutations in p110α protein. Pathogenic/likely pathogenic mutations are shown in red. (E) VAF distribution of *PIK3CA^H1047R^* mutations, and other *PIK3CA* missense mutations classified either as pathogenic/likely pathogenic or uncertain significance/no data available. Medians (red) and quartiles (grey lines) are represented. Two-tailed Mann-Whitney test.

In light of these observations, we set out to investigate the mechanisms by which *Pik3ca^H1047R^* mutant progenitors colonize normal esophagus in mouse esophageal epithelium. The tissue consists of layers of keratinocytes, with progenitor cells residing in the deepest, basal cell layer. Differentiating cells exit the cell cycle and leave the basal layer, migrating towards the epithelial surface where they are shed (Doupe et al., 2012; Piedrafita et al., 2020) (**Figure 2A**). Each progenitor division generates either two progenitor daughters, two non-dividing differentiating cells or one cell of each type (**Figure 2B**) (Doupe et al., 2012; Piedrafita et al., 2020). The probabilities of these outcomes are balanced across the progenitor population so that one progenitor and one differentiating daughter are produced from an average division. This ensures that across the progenitor population, equal numbers of progenitor and differentiating cells are generated, maintaining tissue homeostasis. Mutations may tilt the balance of cell production towards proliferation driving the clonal expansion of mutant progenitors (Alcolea et al., 2014; Fernandez-Antoran et al., 2019; Murai et al., 2018) (**Figure 2B**).

**Figure 2.**
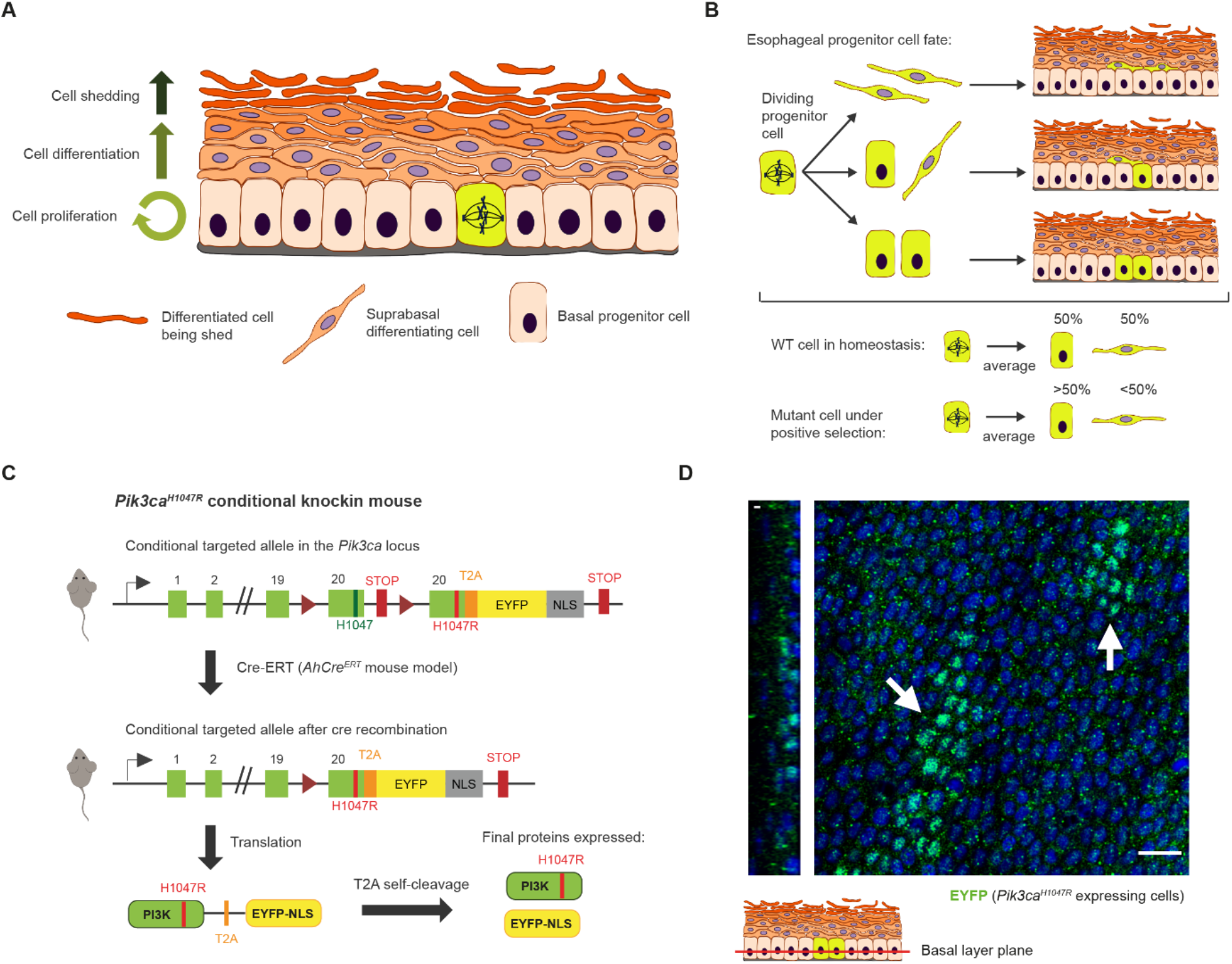
Mouse esophageal epithelium cell behavior and conditional *PIK3CA^H1047R^* transgenic mouse model. (A) Mouse esophageal epithelium. The lowest basal layer contains proliferating progenitor cells and post-mitotic differentiating cells which will enter the suprabasal cell layers. Differentiating cells migrate through the layers to the tissue surface where they are shed. (B) Each progenitor cell division may generate two differentiating cells, two progenitor cells or one of each type. In homeostasis, the probabilities of each outcome are balanced in order to produce equal numbers of progenitor cells and differentiating cells across the epithelium. Mutations under positive selection may tilt the cell fate towards proliferation, producing excess progenitor cells. (C) Schematic illustration of the conditional targeted allele in the *Pik3ca* locus. *Pik3ca* exon 20 was flanked by *loxP* sites (triangles). The engineered duplicate region of *Pik3ca* exon 20 contains the H1047R mutation, and sequences coding for a self-cleaving T2A peptide and an enhanced Yellow Fluorescent Protein (EYFP) followed by a nuclear localization signal (NLS). Prior to Cre-mediated recombination, the wild type p110α protein is expressed; however, after Cre mediated recombination, the allele co-expresses p110α^H1047R^ mutant protein and EYFP-NLS. Cre recombination was mediated by crossing the conditional mutant strain with *AhCre^ERT^* mice which express the Cre recombinase upon treatment with β-naphthoflavone and tamoxifen. (D) Typical confocal z stack image of a region of an esophageal epithelial whole-mount, viewed top down, from an *AhCre^ERT^PIK3CA^H1047R/wt^* mice 3 months after induction with β-naphthoflavone and tamoxifen. An optical section through the basal cell layer is shown, *PIK3CA^H1047R/wt^* clones are indicated by white arrows. YFP immunofluorescence (green), nuclei are stained with DAPI (blue). Scale bar, 20 μm.

To explore the effect of activating *Pik3ca* mutations on progenitor cells we developed a new transgenic mouse model that allowed us to track the fate of individual progenitors induced to express a single allele of the *Pik3ca^H1047R^* mutant in a background of wild type cells. We found that this mutation tilts progenitor cell fate, so that excess of mutant progenitors are produced per average division, a proliferative advantage that promotes mutant clone expansion. We show that *Pik3ca^H1047R^* activates glycolysis via the HIF1α transcription factor in esophageal cells and that levelling the competitive ‘playing field’ with agents that regulate glycolysis blocks the competitive advantage of *Pik3ca^H1047R^* mutant cells *in vitro* and *in vivo*.

## Results

### Generation of inducible *Pik3ca^H1047R-YFP^* knock-in mice

The majority of *Pik3ca* mutant clones found in the normal esophagus carry an activating H1047R mutation in one allele of the *Pik3ca* locus. To determine whether induction of a heterozygous *Pik3ca^H1047R^* allele altered progenitor cell behavior, we developed a new conditional mouse strain, *Pik3ca^fl-H1047R-T2A-YFP-NLS^* (henceforth referred to as *Pik3ca^H1047R-YFP^*). A conditional allele of *Pik3ca^H1047R^*, with a nuclear localized Yellow Fluorescent Protein (YFP) reporter linked to the C-terminus of the *Pik3ca^H1047R^* protein by a T2A self-cleaving peptide (Trichas et al., 2008), was targeted to the *Pik3ca* locus. Following recombination mediated by *Cre* recombinase, the wild type exon 20 of *Pik3ca* is excised and replaced by the mutant exon 20 encoding *Pik3ca^H1047R^* (**Figure 2C**). This design allowed us to track individual *Pik3ca^H1047R-YFP^* cells in a *Pik3ca*^wt/wt^ background, as the recombined mutant cells and their progeny stain positive for YFP (**Figure 2D**). Following T2A cleavage, a 20-amino-acid peptide remained at the C terminus of *Pik3ca^H1047R^* protein. We found that a C-terminally extended p110α^H1047R^ protein could still activate the PI3K/Akt pathway upon transient transfection in NIH3T3 fibroblasts (**Figure S1A**) and also induce modest Akt activation upon *in vitro* recombination of primary esophageal keratinocytes from *Pik3ca^H1047R-YFP^* mice (**Figure S1B-F**).

**Supplementary Figure 1.**
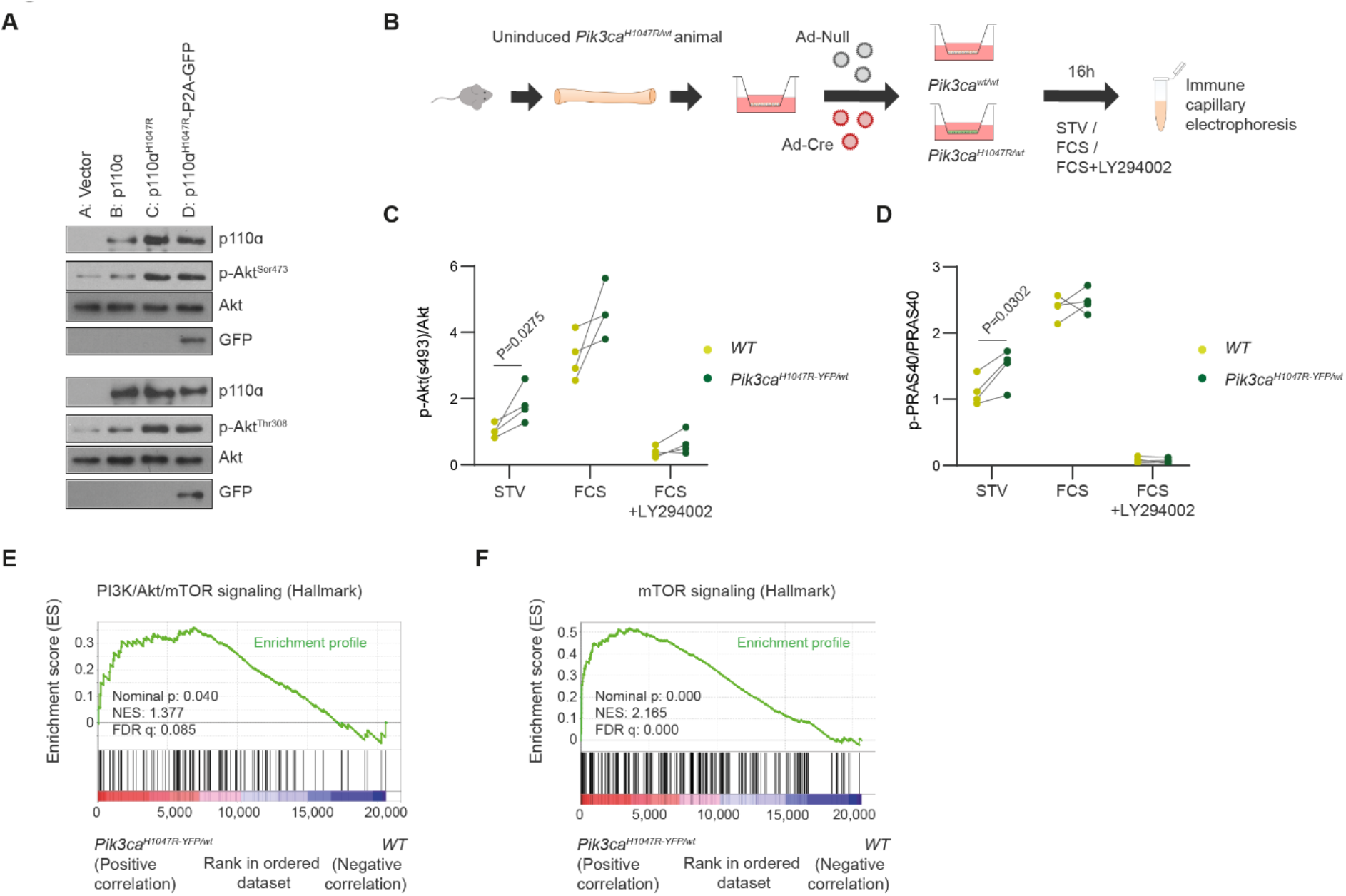
Related to Figure 2. Expression of p110αH1047R mutant protein activates PI3K pathway. (A) Immunoblots of NIH3T3 cells transfected with an empty vector, a vector carrying wild type p110α, p110α^H1047R^ or p110α^H1047R^-P2A-GFP. Cells were starved for 24 h and lysed. Expression of PI3K, GFP, AKT and phosphorylation of AKT on Ser^473^ and Thr^308^ is shown. Images are representative of 3 replicate experiments. (B) Experimental protocol. Primary esophageal keratinocytes were isolated from uninduced *PIK3CA^H1047R-YFP/wt^* mice and infected either with null-adenovirus (uninduced *Pik3ca^wt/wt^* controls) or Cre-adenovirus (induced *PIK3CA^H1047R-YFP/wt^*). Cells were treated in medium without added growth factors and 0.1% serum (Starved, STV) or medium with 20% serum and growth factors (FCS) or FCS plus the PI3K inhibitor LY294002 50 μM. Cells were lysed and protein lysates were analyzed by immune capillary electrophoresis. (C-D) Results from immune capillary electrophoresis of lysates from protocol shown in (B). Total AKT protein and phospho-AKT(Ser473) (C) and total PRAS40 protein and phospho-PRAS40 (D). Two-tailed ratio paired *t*-test. n= 4 biological replicates. (E-F) Gene set enrichment analysis (GSEA) histograms of PI3K/Akt/mTOR and mTOR signaling Hallmark gene sets comparing RNA-seq data from induced *PIK3CA^H1047R-YFP/wt^* and uninduced cells from the same animals maintained in minimal FAD medium. The nominal p-value, the normalized enrichment score (NES) and the false discovery rate (FDR) q-value are indicated. n= 4 independent replicates per condition from one animal each.

### *Pik3ca^H1047R-YFP/wt^* mutant cells outcompete wild type cells in esophageal epithelium

To track the fate of *Pik3ca^H1047R-YFP/wt^* progenitors *in vivo*, we crossed *Pik3ca^H1047R-YFP/wt^* mice with the inducible *Cre* recombinase line *AhCre^ERT^*. *Cre* was induced in adult *AhCre^ERT^Pik3ca^H1047R-YFP/wt^* mice (henceforth termed *Cre-Pik3ca^H1047R-YFP/wt^*) at a level that recombined individual esophageal cells, which went on to generate clones of *Pik3ca* mutant cells (Clayton et al., 2007; Doupe et al., 2012; Kemp et al., 2004). As controls, we induced *AhCre^ERT^Rosa26^flYFP/wt^* (henceforth termed *Cre-RYFP*) animals that are *Pik3ca* wild type and express a neutral YFP reporter after recombination. At multiple time points up to 6 months after induction, animals were culled (**Figure 3A**), the esophagus collected and the entire epithelium isolated, stained and imaged in 3D using confocal microscopy. The number of cells and their location within individual clones was then counted and control and mutant clone size distributions compared (**Table S1)**.

**Figure 3.**
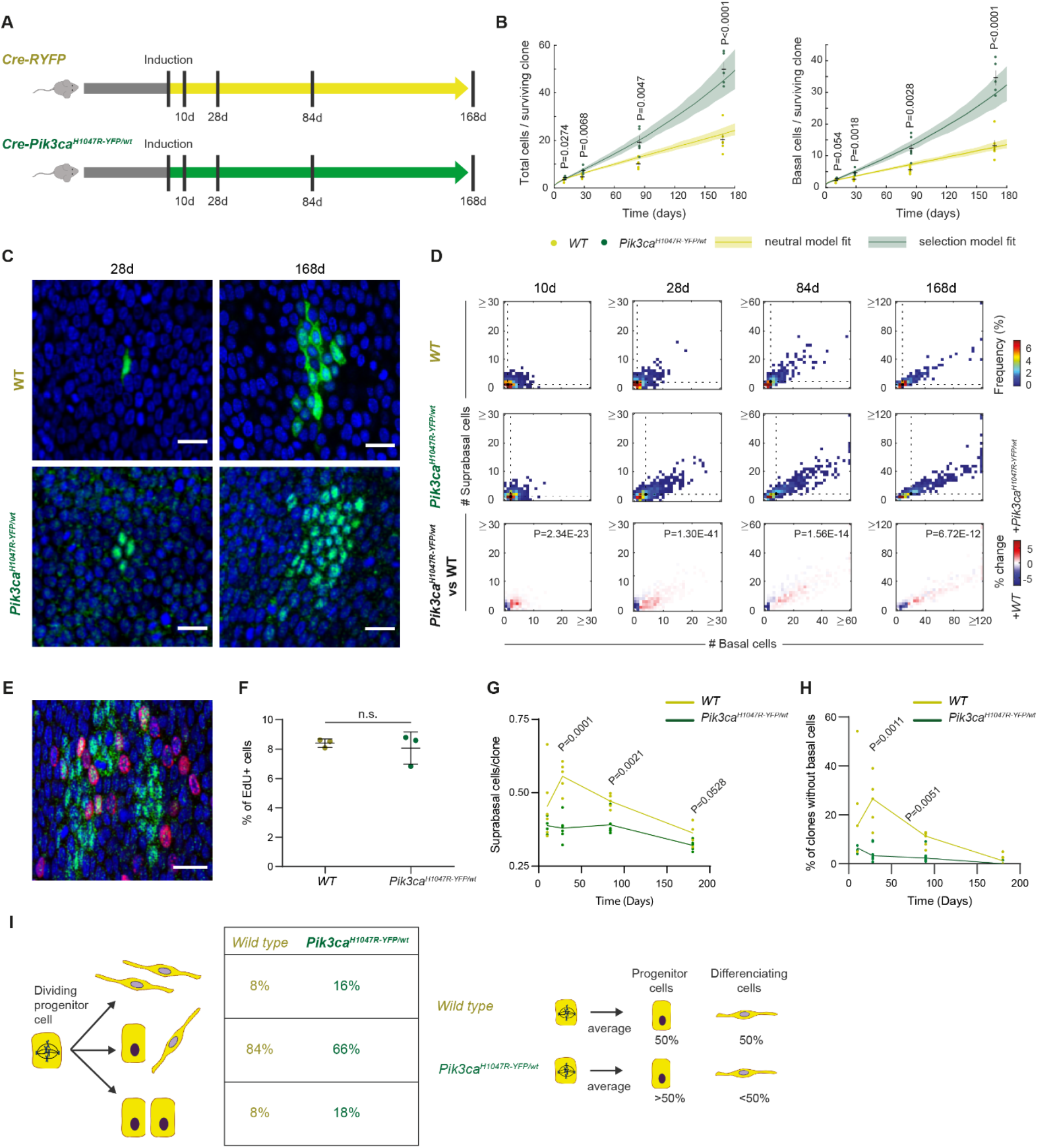
Lineage tracing reveals a bias in *Pik3ca^H1047/wt^* cell fate favoring mutant clone growth. (A) Experimental protocol. *AhCre^ERT^Rosa26^flEYFP/wt^* reporter mice (*Cre-RYFP*) and *AhCre^ERT^Pik3caflH1047R-T2A-YFP-NLS/wt* mice (*Cre-PIK3CA^H1047R-YFP/wt^*) were induced with β-naphthoflavone and tamoxifen and tissue collected at the indicated time points. Wild type (*WT*) clones from *Cre-RYFP* mice and *PIK3CA^H1047R-YFP/wt^* clones from *Cre-PIK3CA^H1047R-YFP/wt^* animals were imaged and cell numbers in each clone quantified. n=317-1522 clones from 4-6 animals per condition (see Table S1 for values). Average total cells per clone (basal layer plus first suprabasal layer cells, left panel) and basal cells per clone (right panel) over time, for all clones with at least one basal cell. Dots indicate the average clone size in a single mouse. Lines and shaded areas represent the best fitting model for the clone size distributions and its plausible intervals (see **Supplementary Text**). Mean and standard error of the mean per condition are indicated in black. Two-tailed unpaired *t*-test. (C) Top down views of confocal images of representative clones (green) for *Cre-RYFP* reporter mice (top panels) and *Cre-PIK3CA^H1047R-YFP/wt^* mice (lower panels) 28 days and 168 days after induction. Nuclei are stained with DAPI (blue). An optical section through the basal cell layer is shown. Scale bars, 20 μm. (D) Heatmaps representing the frequency of clone sizes with the number of basal and first suprabasal cells indicated, for *Cre-RYFP* and *Cre-PIK3CA^H1047R-YFP/wt^* animals. Black dots and dashed lines show geometric median clone size. Lower panels show the differences between *Cre-PIK3CA^H1047R-YFP/wt^* and *Cre-RYFP* animals for each time point. 2D Kolmogorov-Smirnov test. (E) Basal cells (DAPI, blue) of a typical EE whole mount showing *PIK3CA^H1047R-YFP/wt^* cells (green), and EdU^+^ basal cells (red). Scale bar, 20 μm. (F) Percentages of EdU^+^, uninduced (WT) or *PIK3CA^H1047R-YFP/wt^*, basal cells quantified in the same tissues one month after induction. 52,898 cells were quantified from 3 animals including 3,514 *PIK3CA^H1047R-YFP/wt^* cells. Each dot corresponds to an animal. Mean and standard deviation are shown in black. n.s., not significant (Ratio paired *t*-test). (G) Average proportion of suprabasal cells per clone over time, counting basal and first suprabasal cells from clones shown in (D). Each dot corresponds to one animal and the lines connect means. Two-tailed unpaired *t*-test. (H) Proportion of floating clones per mouse. Each dot corresponds to one animal and the lines connect averages. Two tailed unpaired *t*-test. n=4-6 animals per condition (see Table S1 for values). (I) Schematic illustration of *WT* and *PIK3CA^H1047R-YFP/wt^* cell behavior. The model predictions for the proportions of each cell division fate for both genotypes are shown. A bias in the fate of *PIK3CA^H1047R-YFP/wt^* progenitors together with an increased proportion of symmetric divisions results in an increased production of mutant progenitor cells per average cell division in the esophageal epithelium, even with the rate of mutant cell division being the same as that of *WT* cells.

Within 10 days of induction we observed that the total number of cells in *Pik3ca^H1047R-YFP/wt^* clones was larger than that of wild type controls, the difference increasing progressively over 6 months (**Figures 3B and C**). The mutant clones had a higher proportion of basal cells and fewer differentiated (suprabasal) cells compared with controls (**Figure 3D**). These observations indicate *Pik3ca^H1047R-YFP/wt^* cells have a competitive advantage over their wild type neighbors and suggest that mutant progenitors may generate a lower proportion of differentiating and more proliferating progeny than their wild type equivalents.

To test if these results were due to genetic background differences between mutant and control mouse strains, we crossed *Cre-Pik3ca^H1047R-YFP/wt^* mice with the *Rosa26^Confetti/wt^* strain (Snippert et al., 2014) (**Figure S2A**). This triple mutant allows to track *Pik3ca^H1047R-YFP/wt^* expressing clones (labelled with YFP) and non-recombined *Pik3ca* wild type clones, both labelled with red fluorescent protein (RFP) in the same esophagus (**Figures S2B and C**). The results confirmed that the mutant clones expanded more rapidly than wild type clones in the same mouse (**Figures S2D and E**). Therefore, we conclude that *Pik3ca^H1047R-YFP/wt^* progenitors have a competitive advantage over their wild type neighbor cells in the mouse esophageal epithelium.

### *Pik3ca^H1047R^* mutation biases progenitor cell fate towards proliferation

There are several possible cellular mechanisms that may underpin the increased size of *Pik3ca^H1047R^* mutant over neutral, wild type clones. To investigate whether altered cell cycle kinetics in the mutant cells were responsible for this effect, we induced and aged *Cre-Pik3ca^H1047R-YFP/wt^* mice for 1 month. One hour before tissue collection, animals were injected with 5-ethynyl-2′-deoxyuridine (EdU), which labels keratinocytes in S phase of the cell cycle. The percentage of labelled basal cells depends on the duration of S phase/cell cycle time and the proportion of basal layer cells that are progenitors. The proportion of EdU^+^ basal cells was similar in mutant cells compared with wild type cells in the same esophagus (**Figures 3E and F**). These results argue that the cell-cycle time and the proportion of cycling progenitors are not substantially altered by the *Pik3ca^H1047R^* mutation.

**Supplementary Figure 2.**
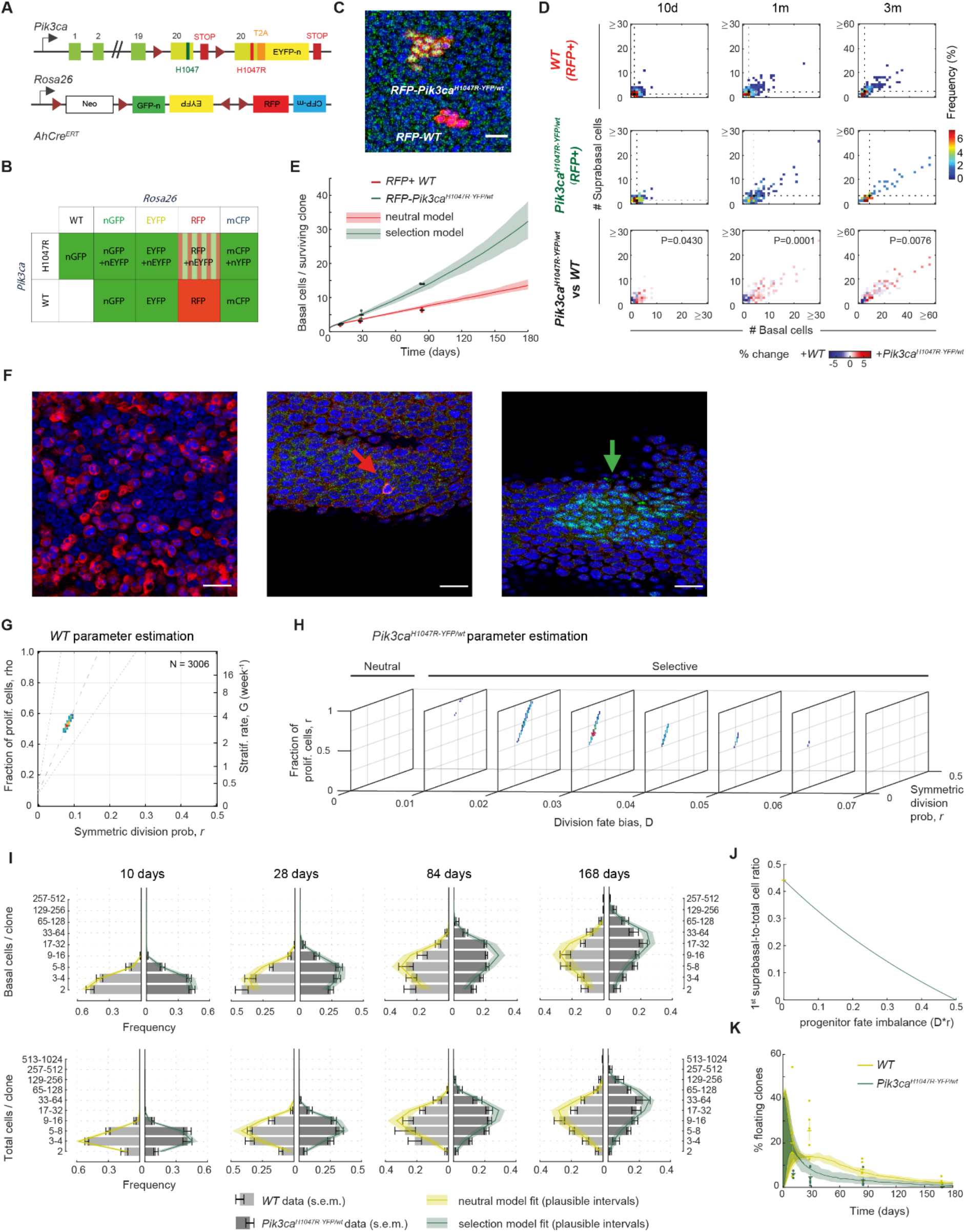
Related to Figure 3. Lineage tracing in *Pik3ca^H1047R-YFP/wt^Rosa26^confetti/wt^AhCre^ERT^* mice, apoptosis measurement and mathematical modeling for *Cre-RYFP* and *Cre-Pik3ca^H1047R-YFP/wt^* lineage tracing. (A) Multicolor lineage tracing in *PIK3CA^H1047R-YFP/wt^Rosa26^confetti/wt^AhCre^ERT^* (*Cre-PIK3CA^H1047R-YFP/wt^-Confetti*) animals. The confetti reporter allele encodes four fluorescent proteins. After random Cre-mediated recombination induced by treating with β-naphthoflavone and tamoxifen, one fluorescent reporter is expressed by the cell and its progeny (clone). Some fluorescently labelled clones will also recombine *Pik3ca* locus expressing the mutation together with an EYFP reporter. (B) Table showing the possible color combinations obtained from (A) after immunofluorescence against GFP to detect *PIK3CA^H1047R-YFP/wt^*. GFP antibody recognizes GFP, EYFP and CFP together with EYFP expressed from the *Pik3ca* locus. Only RFP+ clones wild type or *PIK3CA^H1047R-YFP/wt^* can be distinguished and quantified. (C) Top down views of confocal images of an esophageal epithelium whole-mount of a *Cre-PIK3CA^H1047R-YFP/wt^-Confetti* animal 84 days after induction, showing a RFP^+^ *Pik3ca^wt/wt^* and a RFP^+^ *PIK3CA^H1047R-YFP/wt^* clone. Nuclei stained with DAPI (blue). Scale bar, 20 μm. (D) Heatmaps representing the frequency of clone sizes, with the number of basal and first suprabasal cells indicated, of RFP^+^ clones observed in *Cre-PIK3CA^H1047R-YFP/wt^-Confetti* animals. RFP^+^ clones from each animal were classified into *Pik3ca^wt/wt^* and *PIK3CA^H1047R-YFP/wt^* by immunofluorescence. Black dots and dashed lines show geometric median clone size. Lower panels show the differences between *Pik3ca^wt/wt^* and *PIK3CA^H1047R-YFP/wt^* clones. 2D Kolmogorov-Smirnov test. n=255-469 clones in total from 2-3 animals per condition (see Table S1 for values). (E) Average basal cells per clone over time, considering all clones with at least one basal cell. Dots indicate the average clone size of a mouse. Mean and standard error of the mean per condition are indicated in black. Lines and shaded areas represent the best fitting model for the clone size distributions shown in **Figure 3** and its plausible intervals (see **Supplementary Text**). (F) Confocal images of an esophageal epithelium basal layer whole-mount stained for activated Caspase 3 (red), GFP (*PIK3CA^H1047R-YFP/wt^* cells, green) and DAPI (blue). Left panel shows a UV irradiated sample as a positive control, exposed to ultraviolet radiation and maintained as explant culture. Middle and right panels are images from the same tissue showing an apoptotic cell (red arrow) and a *PIK3CA^H1047R-YFP/wt^* clone (green arrow). Scale bars, 20 μm. (G-H) Results of parameter inference for wild type (G) and mutant (H) progenitor cells. Distributions of the number of basal cells/clone were fitted (see **Supplementary Text**) and heatmaps show most-likely parameter values according to likelihood inference for a neutral single-progenitor model with balanced fates (wild type, G) and when extending the analysis to a single-progenitor model with imbalanced fates, displaying selection (mutant, H). In the latter case, different values for the parameter of division fate bias Δ were considered (Δ =0 means neutral behavior). Red asterisk: maximum likelihood estimate (MLE). Colored regions fall within 95% CI (uncolored regions are out of bounds). (I) Distributions of wild type (light grey) and mutant (dark grey) clone sizes. Number of basal cells/clone, and number of total (basal + first suprabasal) cells/clone are displayed (top and bottom panels, respectively) (sizes grouped in powers of two). Error bars: experimental mean ± s.e.m. Overlaid are MLE model fits (shaded areas represent 95% plausible intervals given the total number of clones counted at each time point). (J) Theoretical prediction for the effect of a progenitor fate imbalance on the relative proportion of first suprabasal cells. The first suprabasal-to-total cell ratio decreases with Δ*r following a rational decay function (see **Supplementary Text**) (initial departure point corresponds to wild type value). (K) Proportion of floating clones (i.e. suprabasal clones having no basal attachment) over time. Yellow and green dots correspond to values in individual wild type and mutant mice, respectively (error bars: mean ± s.e.m.). Overlaid are MLE model fits once changes in suprabasal-to-total cell ratio over time were considered (shaded areas defined as in I).

Another potential mechanism of cell competition is by promoting apoptosis of neighboring cells (de la Cova et al., 2014). However, there was a negligible level of apoptosis in wild type cells, whether adjacent to or distant from mutant clones in induced *Cre-Pik3ca^H1047R-YFP/wt^* animals (**Figure S2F**).

Finally, we investigated whether *Pik3ca^H1047R-YFP/wt^* clone behavior could be explained by altered progenitor cell fate. Even when cell division rates are similar, mutant populations could still expand by producing more progenitor than differentiating daughter cells per average cell division, as previously observed with some other genetic mutants (Alcolea et al., 2014; Murai et al., 2018; Piedrafita et al., 2020). Mathematical modeling revealed that wild type clones follow a neutral model of cell competition, as described previously, with equal proportions of proliferating and differentiating cells produced from the average cell division (**Figures 3I and S2G**) (Doupe et al., 2012; Piedrafita et al., 2020). However, *Pik3ca^H1047R-YFP/wt^* clone dynamics is explained by a non-neutral model where mutant progenitors have altered division outcome probabilities (**Supplementary text**). There is a fate bias towards proliferation, so that, over the mutant progenitor population, the average cell division generates an excess of proliferating over differentiating progeny, explaining the clonal growth advantage over wild type cells (**Figure 3I and S2H**). This simple model fits both the observed basal cell and total (basal plus suprabasal) clone size distributions and averages (**Figures 3B and S2I**). The model also predicts that a progenitor fate imbalance should result in a decreased proportion of suprabasal cells per clone (**Figure S2J**) and a reduction in the number of fully differentiated clones lacking any basal cells (floating clones) in the mutant (**Figure S2K, Supplementary Text**). These predictions also fit with experimental data since both the proportion of suprabasal cells per clone and floating clones were significantly reduced in the mutant (**Figures 3G and H**).

Therefore, *Pik3ca^H1047R-YFP/wt^* keratinocytes show a bias in basal cell fate towards the generation of more progenitor than differentiating daughters, resulting in mutant cells having a competitive advantage over wild type cells in the esophageal epithelium (**Figure 3I, Supplementary video**)

### *Pik3ca^H1047R-YFP/wt^* mutant cell fitness depends on the level of PI3K pathway activation in neighboring wild type cells

To further investigate the basis of the mutant cell advantage over wild type cells we used a 3D stratified primary culture system suitable for long-term cell competition studies. We generated primary esophageal keratinocyte cultures from *Rosa^RYFP/RYFP^* (*Pik3ca^wt/wt^*, henceforth referred to as *WT-RYFP*) and *Pik3ca^H1047R-YFP/wt^* mice, and induced recombination by infecting these cultures with adenovirus encoding *Cre* recombinase (**Figures S3A and B**). Due to the much higher level of YFP reporter expression in *WT-RYFP* compared with *Pik3ca^H1047R-YFP/wt^* cells, they can be easily identified by flow cytometry (**Figure S3C**).

**Supplementary Figure 3.**
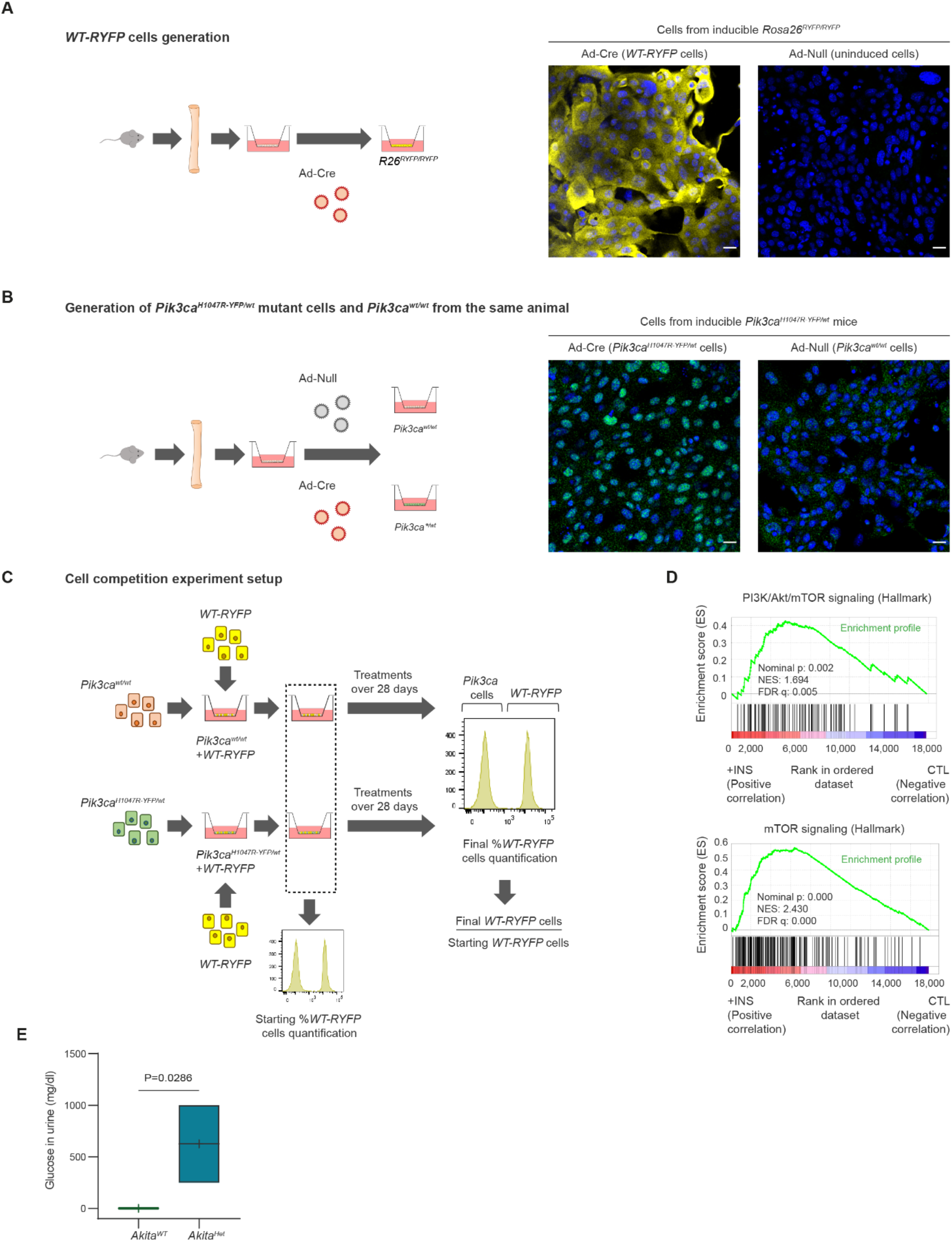
Related to Figure 4. Set up for cell competition experiments. (A-B) Generation of fully induced primary *WT-RYFP* (A) and *PIK3CA^H1047R-YFP/wt^* (B) esophageal keratinocytes. Primary esophageal keratinocytes were isolated from uninduced *Rosa26^RYFP/RYFP^* animals (A) or uninduced *PIK3CA^H1047R-YFP/wt^* animals (B). Cells were incubated either with Cre-expressing adenovirus (Ad-Cre) or null adenovirus (Ad-Null). Right panels show a representative image of an immunofluorescence against YFP (green) and DAPI (blue) of Ad-Cre or Ad-Null treated cultures. Scale bars, 20 μm. (C) Cell competition experimental protocol. *PIK3CA^H1047R-YFP/wt^* and *Pik3ca^wt/wt^* primary keratinocytes obtained in (B) were mixed with *WT-RYFP* cells obtained in (A). Upon reaching confluence, cultures were changed to minimal FAD medium or the specified treatment. Cells were collected at the start and end of the treatment and the proportion of *WT-RYFP* cells was quantified by flow cytometry. The proportion of *WT-RYFP* after treatment was normalized to the initial *WT-RYFP* cell proportion. (D) GSEA histograms of PI3K/Akt/mTOR and mTOR signalling Hallmark gene sets comparing RNA-seq data from control (CTL) and 5 μg/ml insulin (+INS) treated *PIK3CA^H1047R-YFP/wt^* cells from the same animals. The nominal p-value, the normalized enrichment score (NES) and the false discovery rate (FDR) q-value are indicated. n= 4 independent replicates per condition from one animal each. (E) Glucose levels in urine of the mice used in Figure 4 I-K at the day of induction. Two-tailed Mann-Whitney test. n=4 animals per condition.

For the cell competition studies, we mixed *WT-RYFP* keratinocytes with either induced or uninduced *Pik3ca^H1047R-YFP/wt^* cells and followed how the proportion of *WT-RYFP* cells changed over time (**Figures 4A and S3C**). We first established that when uninduced *Pik3ca^H1047R-YFP/wt^* cells were mixed with *WT-RYFP* cells, the proportion of cells of both strains remained constant over time, meaning their competition is neutral (**Figure 4B upper panels and C**). In addition, the ratio of suprabasal:basal cells was similar in both subpopulations after 14 days of cell competition (**Figure 4D**). However, when induced *Pik3ca^H1047R-YFP/wt^* and *WT-RYFP* cells were co-cultured, the *Pik3ca^H1047R-YFP/wt^* cells almost completely took over the culture within 28 days (**Figure 4B lower panels and C**). Moreover, the suprabasal:basal ratio of induced *Pik3ca^H1047R-YFP/wt^* cells was lower than for the *WT-RYFP* cells in the same culture (**Figure 4D**). We conclude that *Pik3ca^H1047R-YFP/wt^* mutant cells retain their competitive advantage over wild type cells *in vitro*.

**Figure 4.**
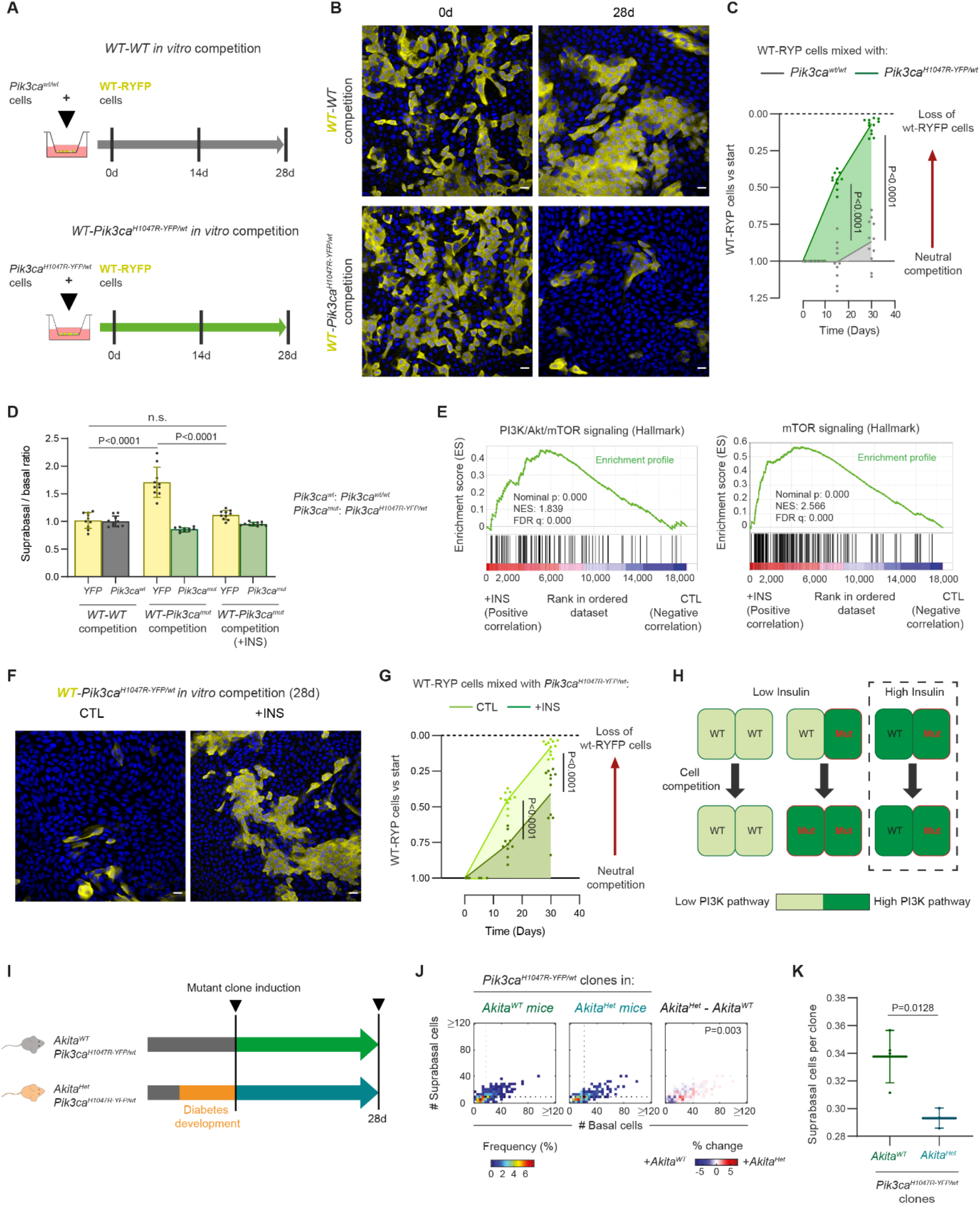
Differential PI3K pathway activation modulates the competitive advantage *of PIK3CA^H1047R-YFP/wt^* cells. (A) Experimental protocol for *in vitro* competition. *In vitro* induced *Rosa26^flYFP/flYFP^* cells (*WT-RYFP*) were mixed with *in vitro* induced *PIK3CA^H1047R-YFP/wt^* cells or uninduced *Pik3ca^wt/wt^* cells from the same animals (see **Figure S3**). Once a confluent culture was achieved, cells were kept for 28 days in culture with minimal FAD medium, or the specified treatment, for the duration of the experiment. Samples were collected at the start of the treatment and at 14 and 28 days. (B) Confocal z stack image representative of the specified mixed culture and time of treatment. An optical section through the basal cell layer is shown. YFP immunofluorescence (yellow), nuclei are stained with DAPI (blue). Scale bar, 20 μm. (C) Quantification by flow cytometry of the proportion of *WT-RYFP* cells *versus* the start of the experiment at the specified time points. Each dot represents a primary culture from a different animal. n=10-11 primary cultures from individual animals per condition. Two-tailed ratio paired *t*-test. (D) Proportion of suprabasal *versus* basal cells of each subpopulation in mixed cultures 14 days after the start of the experiment. +INS indicate cultures treated with 5 μg/ml insulin. n=10 primary cultures from individual animals per condition. n.s., not significant. Two-tailed unpaired *t*-test. (E) GSEA histograms of PI3K/Akt/mTOR and mTOR signalling Hallmark gene sets comparing RNA-seq data from control (CTL) and 5 μg/ml insulin (+INS) treated wild type cells from the same animals. The nominal p-value, the normalized enrichment score (NES) and the false discovery rate (FDR) q-value are indicated. n= 4 independent replicates per condition from one animal each. (F) Confocal z stack image representative of the specified mixed culture and condition after 28 days of continuous treatment. An optical section through the basal cell layer is shown. YFP immunofluorescence (yellow), nuclei are stained with DAPI (blue). Scale bar, 20 μm. (G) Quantification by flow cytometry of the proportion of *WT-RYFP* cells mixed with *PIK3CA^H1047R-YFP/wt^* cells, at the specified time points *versus* the start of the experiment. Cells were treated either in minimal FAD medium or treated with 5 μg/ml insulin (+INS) for the duration of the experiment. Each dot represents a primary culture from an animal. n=10-11 primary cultures from individual animals per condition. Two-tailed ratio paired *t*-test. (H) Summary of the results representing *PIK3CA^H1047R-YFP/wt^* (Mut) *versus* wild type (WT) competition in control (Low insulin) and insulin (High Insulin) conditions; in relation to the PI3K pathway activation in each subpopulation. (I) Experimental protocol: *Cre-PIK3CA^H1047R-YFP/wt^* mice were bred into *Ins2^Akita/wt^* (*Akita^Het^*) mice obtaining *Cre-PIK3CA^H1047R-YFP/wt^-Akita^Het^* and *Cre-PIK3CA^H1047R-YFP/wt^-Akita^WT^* littermates. After diabetes development in *Akita^Het^* mice, they were induced with β-naphthoflavone and tamoxifen and collected after 28 days. (J) Heatmaps representing the frequency of *PIK3CA^H1047R-YFP/wt^* clone sizes, with the number of basal and first suprabasal cells indicated, observed in *Cre-PIK3CA^H1047R-YFP/wt^-Akita^Het^* and *Cre-PIK3CA^H1047R-YFP/wt^-Akita^WT^* littermates (left panels). Heatmap showing the differences in *PIK3CA^H1047R-YFP/wt^* clone sizes between *Cre-PIK3CA^H1047R-YFP/wt^-Akita^Het^* and *Cre-PIK3CA^H1047R-YFP/wt^-Akita^WT^*. n=257 and 402 clones respectively, from 4 animals per condition (see Table S1 for numbers). 2D Kolmogorov-Smirnov test. (K) Average proportion of suprabasal cells in *PIK3CA^H1047R-YFP/wt^* clones in *Cre-PIK3CA^H1047R-YFP/wt^-Akita^Het^* and *Cre-PIK3CA^H1047R-YFP/wt^-Akita^WT^* littermates. Each dot corresponds to one animal. Bars show mean and standard deviation. Two-tailed unpaired *t*-test.

If the advantage of *Pik3ca^H1047R-YFP/wt^* mutation depends on over-activation of the PI3K/mTOR axis, we reasoned that activating this pathway in wild type cells would decrease the fitness advantage of the mutant cells, by levelling out the signaling differences between the two genotypes. High concentrations of insulin activate PI3K/mTOR via the insulin and IGF1 receptors (Boucher et al., 2010). Thus, in a mixed culture both wild type and mutant cells might experience strong PI3K/mTOR activation. We treated mixed cultures with a dose of insulin that induced transcriptional changes consistent with PI3K/mTOR pathway activation in both wild type and mutant cells (**Figure 4E and S3D**). The advantage of the *Pik3ca^H1047R-YFP/wt^* over *Pik3ca^wt/wt^* cells was substantially reduced by insulin treatment (**Figure 4F and G**). In addition, insulin treatment lowered the ratio of suprabasal:basal compartment in wild type cells close to that seen when competing with uninduced *Pik3ca*^wt/wt^ cells (**Figure 4D**). We conclude that differential activation of the PI3K pathway underpins the competitive advantage of mutant over wild type cells (**Figure 4H**).

These results suggest that insulin levels may alter the competitiveness of *Pik3ca^H1047R-YFP/wt^* clones *in vivo*. To explore this, we turned to the Akita mouse model of type-1 diabetes, which harbors a mutation in the insulin-2 gene that results in reduced circulating insulin levels as mice age (Oyadomari et al., 2002; Yoshioka et al., 1997). We bred *Cre-Pik3ca^H1047R-YFP/wt^* mice onto an *Akita^Het^* (diabetic) or *Akita^wt^* (non-diabetic) background. Clonal recombination was induced after the onset of the diabetes in *Akita^Het^* mice (**Figure S3E**) and clones analyzed one month after induction (**Figure 4I**). *Pik3ca^H1047R-YFP/wt^* clones showed a reduced proportion of differentiated cells per clone in the diabetic background compared to non-diabetic littermates (**Figures 4J and K**), indicating that the fitness *Pik3ca^H1047R-YFP/wt^* mutant relative to wild type cells is higher when insulin levels in blood are low. Therefore, both *in vitro* and *in vivo*, insulin levels modulate *Pik3ca^H1047R-YFP/wt^* cell competition with wild type cells.

### *Pik3ca^H1047R^* mutation activates HIF1α and aerobic glycolysis

The results above argue that mutant cells may have a differential activation of the pathways downstream of PI3K to gain their competitive advantage over wild type cells. To investigate this, we compared the gene expression of induced and uninduced *Pik3ca^H1047R-YFP/wt^* cultures generated from the same mice. RNA sequencing revealed 301 upregulated and 195 downregulated transcripts (adjusted p-value<0.05) in the mutant cells (**Figures 5A and B**). 47% of the upregulated genes (transcripts with an adjusted p-value<0.01) were known or predicted direct targets of the HIF1α transcription factor (**Figure 5C**), a downstream effector of the PI3K/mTOR pathway (Denko, 2008; Rohwer et al., 2019; Xie et al., 2019). Consistent with activation of the PI3K/mTOR/HIF1 axis by the *Pik3ca^H1047R^* mutation, GSEA and KEGG pathway analysis showed an enrichment of the Hypoxia gene set and the HIF1α signaling pathway (**Figures 5D and E**). HIF1α switches cell metabolism from mitochondrial oxidative phosphorylation towards aerobic glycolysis, the metabolic conversion of glucose to lactate in the presence of oxygen to produce energy (Denko, 2008). Consistent with this function of HIF1α, gene expression analysis showed a significant upregulation of genes encoding for all glycolysis pathway enzymes in *Pik3ca^H1047R^* cells (**Figures 5E, F and G**). In the mutant cells, the expression of HIF1α target genes that promote a metabolic switch to aerobic glycolysis (*Higd1a, Bhlhe40, Bnip3, Pdk1, Ndufa4l2 and Pfkbf3*) (Ameri et al., 2015; Chang et al., 2019; Kim et al., 2006; Rikka et al., 2011; Tello et al., 2011; Yi et al., 2019) and genes that promote the export of lactate and protons to reduce the intracellular acidification derived from a glycolytic metabolism, was also increased (Dovmark et al., 2017; Mboge et al., 2018) (**Figures 5F and G**). To confirm the glycolytic switch in mutant cells we used high resolution respirometry which allows the measurement of the oxygen consumption rate (OCR, proportional to mitochondrial oxidation) and extracellular acidification rate (ECAR, proportional to the glycolysis to lactate) (**Figure 5F**). The ratio of OCR and ECAR indicates whether cells are more oxidative or more glycolytic (Zhang et al., 2012). *Pik3ca^H1047R-YFP/wt^* keratinocytes have a significantly reduced OCR/ECAR ratio, confirming a shift to aerobic glycolysis (**Figure 5H**). We conclude that HIF1α is activated in *Pik3ca^H1047R-YFP/wt^* cells and drives a switch to aerobic glycolysis (**Figure 5I**).

**Figure 5.**
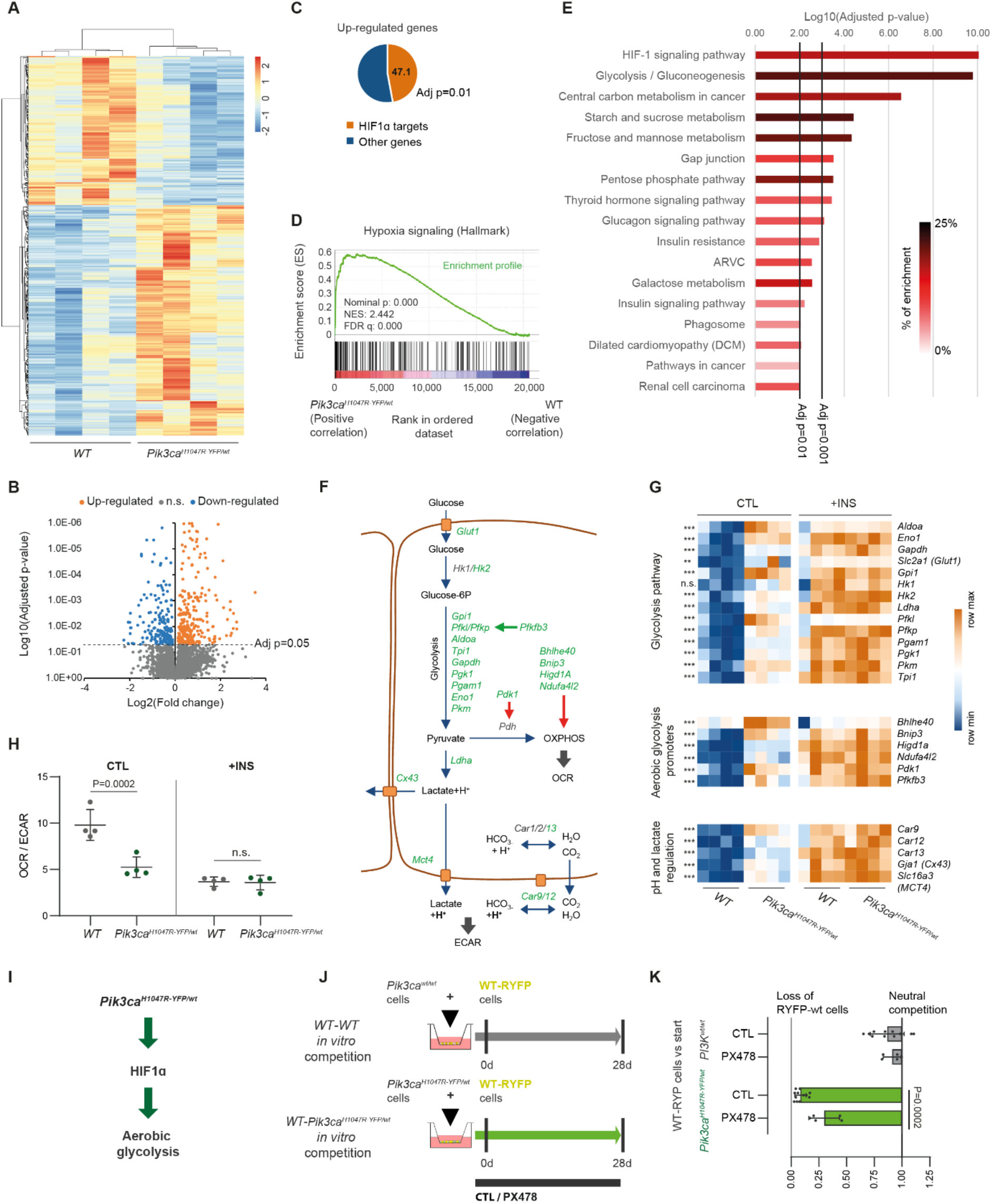
*PIK3CA^H1047R-YFP/wt^* primary esophageal keratinocytes activate HIF1α transcription targets and glycolysis. RNA-seq analysis of gene expression differences between *in vitro* induced *PIK3CA^H1047R-YFP/wt^* and uninduced *Pik3ca^wt/wt^* primary esophageal keratinocytes maintained in minimal FAD medium. Biological quadruplicates were analysed, each sample from a different animal, with an induced and uninduced pair of samples per animal. Adjusted p-values from Wald test corrected for multiple testing using the Benjamini and Hochberg method are shown. (A-B) Heatmap (A) and Volcano plot (B) showing the up-regulated and down-regulated transcripts (adjusted p<0.05) comparing induced *PIK3CA^H1047R-YFP/wt^* and uninduced *Pik3ca^wt/wt^* cells from the same animals. (C) Proportion of HIF1α target genes among the genes significantly up-regulated (adjusted p<0.01) in *PIK3CA^H1047R-YFP/wt^* cells. (D) GSEA histograms of the Hypoxia Hallmarks gene set, comparing *PIK3CA^H1047R-YFP/wt^* cells *versus* uninduced *Pik3ca^wt/wt^* cells. The nominal p-value, the normalized enrichment score (NES) and the false discovery rate (FDR) q-value are indicated. (E) KEGG pathway enrichment analysis for up-regulated genes (adjusted p<0.01) between *PIK3CA^H1047R-YFP/wt^* cells and uninduced *Pik3ca^wt/wt^* cells. Pathways are ordered according to its adjusted p-value and the intensity of the bar color shows its enrichment in the dataset. (F) Scheme of the glycolysis pathway showing the genes implicated and some pathway regulators together with proteins that regulate the internal pH homeostasis. Genes in green are up-regulated in *PIK3CA^H1047R-YFP/wt^* primary keratinocytes. Green and Red arrows show activating and inhibiting processes respectively. OXPHOS: Oxidative Phosphorylation. Oxygen Consumption Ratio (OCR) is proportional to the use of glucose through OXPHOS. Extracellular Acidification Ratio (ECAR) is produced by the protons exported to the extracellular space during aerobic glycolysis. (G) Heatmaps comparing uninduced *Pik3ca^wt/wt^* (WT) and *PIK3CA^H1047R-YFP/wt^* primary keratinocytes in control (CTL) or treated with 5 μg/ml insulin (+INS). Heatmaps show glycolysis pathway genes, genes regulating aerobic glycolysis, and genes regulating the intracellular pH and lactate transport. Statistical tests are performed between CTL samples. None of those genes was differentially expressed between *Pik3ca^wt/wt^* and *PIK3CA^H1047R-YFP/wt^* in the +INS condition. ***p<0.001 and n.s., not significant. Wald test corrected for multiple testing using the Benjamini and Hochberg method. (H) Basal OCR to ECAR ratios of *Pik3ca^wt/wt^* (WT) and *PIK3CA^H1047R-YFP/wt^* primary keratinocytes in control (CTL) or treated with 5 μg/ml insulin (+INS). Basal OCR and ECAR were assessed using the Seahorse Extracellular Flux Analyser. Each dot represents primary cells obtained from one animal (n=4 animals) using 4-5 technical replicates per animal. OCR/ECAR ratios are presented as average and standard deviation. n.s., not significant. Two-tailed ratio paired *t*-test. (I) Model showing how *PIK3CA^H1047R-YFP/wt^* cells activate HIF1α which in turn activates the aerobic glycolysis through its target genes. (J) *In vitro* cell competition assay. *In vitro* induced *Rosa26^flYFP/flYFP^* primary esophageal keratinocytes (*WT-RYFP*) were mixed with *in vitro* induced *PIK3CA^H1047R-YFP/wt^* cells or uninduced controls from the same animals. Once a confluent culture is achieved, cells were treated for 28 days in culture with minimal FAD medium +/-PX-478 (10 μM). Samples were collected at the start of the treatment and at 28 days and the proportion of *WT-RYFP* at the end of the experiment *versus* at the beginning was calculated. (K) Quantification by flow cytometry of the proportion of *WT-RYFP* cells from (J). Each dot represents a primary culture from an animal and mean and standard deviation are shown. n=5-11. Two tailed unpaired *t*-test.

These results suggest HIF1α may be a key effector of the *Pik3ca^H1047R-YFP/wt^* cell phenotype. To test this, we treated mixed cultures of induced *Pik3ca^H1047R-YFP/wt^* and *WT-RYFP* cells with the HIF1α inhibitor PX478 (Welsh et al., 2004) (**Figure 5J**). The advantage of mutant over wild type cells was significantly reduced in the presence of this inhibitor (**Figure 5K**), arguing that activation of HIF1α contributes to the competitive advantage of *Pik3ca^H1047R-YFP/wt^* mutant cells.

We showed above that treatment with a high dose of insulin reduced the mutant cell advantage by causing PI3K pathway over-activation in wild type and mutant cells (**Figure 4E-G and S3D**). We speculated that this treatment may act via HIF1α and glycolysis activation. Transcriptional analysis showed that 82% of the genes upregulated in the *Pik3ca^H1047R-YFP/wt^* mutant cells were also induced in wild type cells upon insulin treatment (**Figure S4A and B**). Insulin treatment abrogated the differences in gene expression between wild type and mutant cells (**Figures S4C**), particularly in HIF1α targets (**Figures S4D**) and glycolysis-related genes (**Figures 5G**).

Consequently, the OCR/ECAR ratio was similar in mutant and wild type cells upon insulin treatment (**Figure 5H**). These observations suggest that insulin treatment levels out the competitive imbalance between wild type and mutant cells by over-activating the PI3K/HIF1α/glycolysis axis both in wild type and mutant cells, supporting the hypothesis that a switch to glycolysis accounts for the increased fitness of mutant over wild type cells.

**Supplementary Figure 4.**
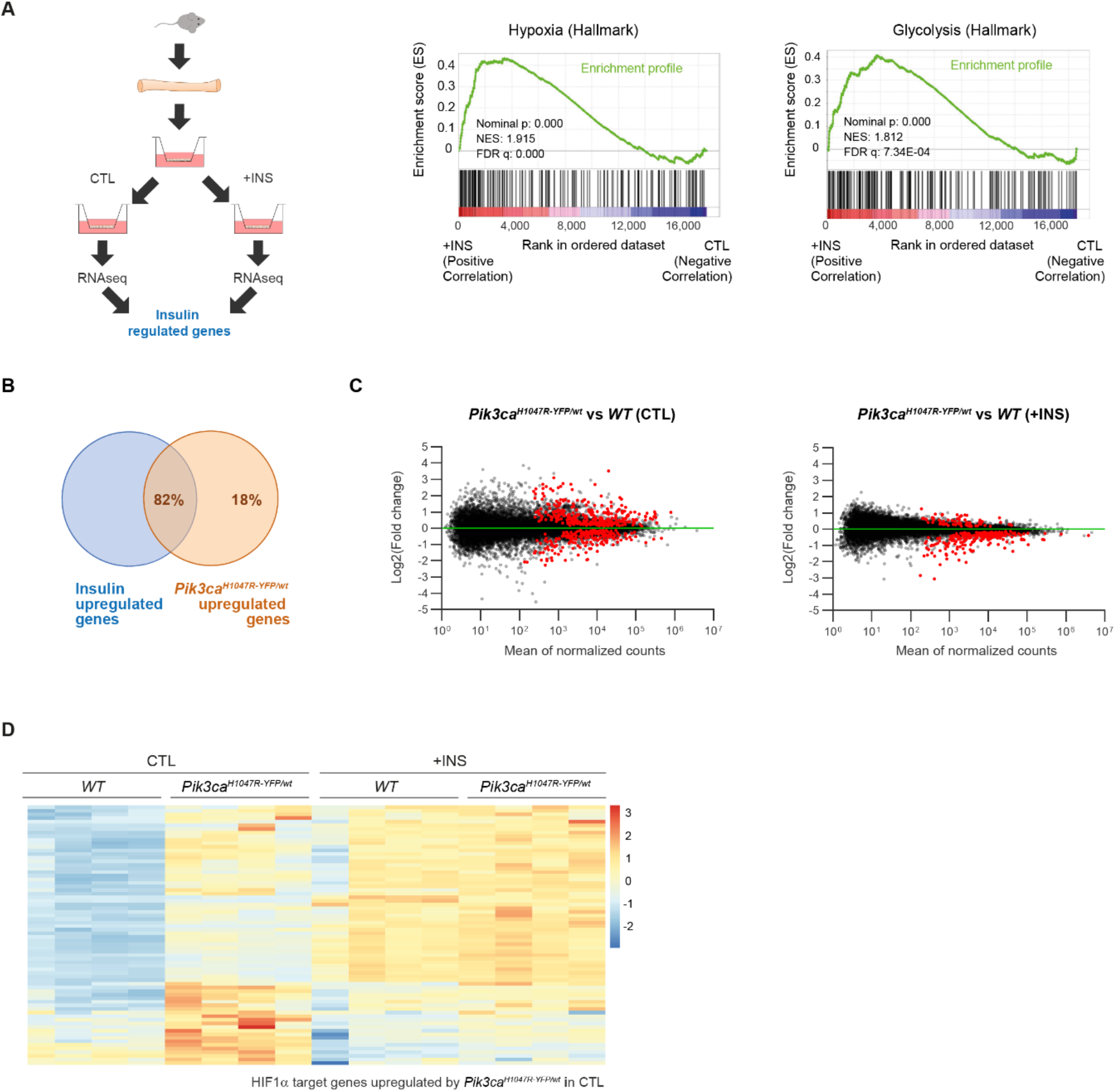
Related to Figure 5. Insulin treatment activates HIF1α and Glycolysis pathways reducing most differences generated by *PIK3CA^H1047R^* mutation. RNA-seq analysis of gene expression differences between *in vitro* induced *PIK3CA^H1047R-YFP/wt^* and uninduced *Pik3ca^wt/wt^* primary esophageal keratinocytes maintained in minimal FAD medium (CTL) or treated with 5 μg/ml insulin (+INS). Biological quadruplicates were analyzed, each sample from a different animal, with an induced and uninduced pair of samples per animal. Adjusted p-values from Wald test corrected for multiple testing using the Benjamini and Hochberg method are shown. (A) RNA-seq analysis of culture of wild type primary esophageal keratinocytes cultured in minimal FAD medium alone (CTL) or with 5 μg/ml insulin (INS). GSEA histograms of the Hypoxia and Glycolysis Hallmarks gene sets, in CTL *versus* INS samples. The nominal p-value, the normalized enrichment score (NES) and the false discovery rate (FDR) q-value are indicated. (B) Venn diagram showing the proportion of genes up-regulated in the *PIK3CA^H1047R-YFP/wt^* cells which are also up-regulated by the insulin treatment in wild type cells. (C) MA plots of RNA-seq data of cultures from induced *PIK3CA^H1047R-YFP/wt^* and *Pik3ca^wt/wt^* (WT) uninduced primary esophageal keratinocytes comparing CTL and +INS treatments; red indicates differentially expressed transcripts with adjusted p<0.05. (D) Heatmap showing the HIF1α target genes significantly up-regulated (adjusted p<0.01) in *PIK3CA^H1047R-YFP/wt^* cells in the CTL condition. Heatmap shows *PIK3CA^H1047R-YFP/wt^* and WT uninduced primary esophageal keratinocytes comparing CTL and +INS treatments.

### Metformin and DCA neutralize the competitive advantage of *Pik3ca^H1047R-YFP/wt^* clones

To test the metabolic dependency of the competitive advantage of *Pik3ca^H1047R-YFP/wt^* cells, we investigated two complementary approaches to reduce the imbalance in glycolysis between mutant and wild type cells. We tested metformin (MET), a widely used antidiabetic agent that enhances aerobic glycolysis (Martin-Montalvo et al., 2013) and dichloroacetate (DCA), a non-selective agent that inhibits pyruvate dehydrogenase kinase-1 (PDK1), hereby forcing glycolysis-derived pyruvate to be oxidized in the mitochondria instead of being transformed into lactate, thus favoring glucose oxidation at the expense of aerobic glycolysis (Michelakis et al., 2008) (**Figures 6A and S5A**). Mixed cultures of *Pik3ca^H1047R-YFP/wt^* and *WT-RYFP* cells were treated with either MET or DCA (**Figure 6B**). Both agents reduced the expansion of *Pik3ca^H1047R-YFP/wt^* cells *in vitro* suggesting that the competitive advantage of *Pik3ca^H1047R-YFP/wt^* mutant cells is attenuated by reducing metabolic differences between mutant and wild type cells (**Figures 6 C, D and E**).

**Figure 6.**
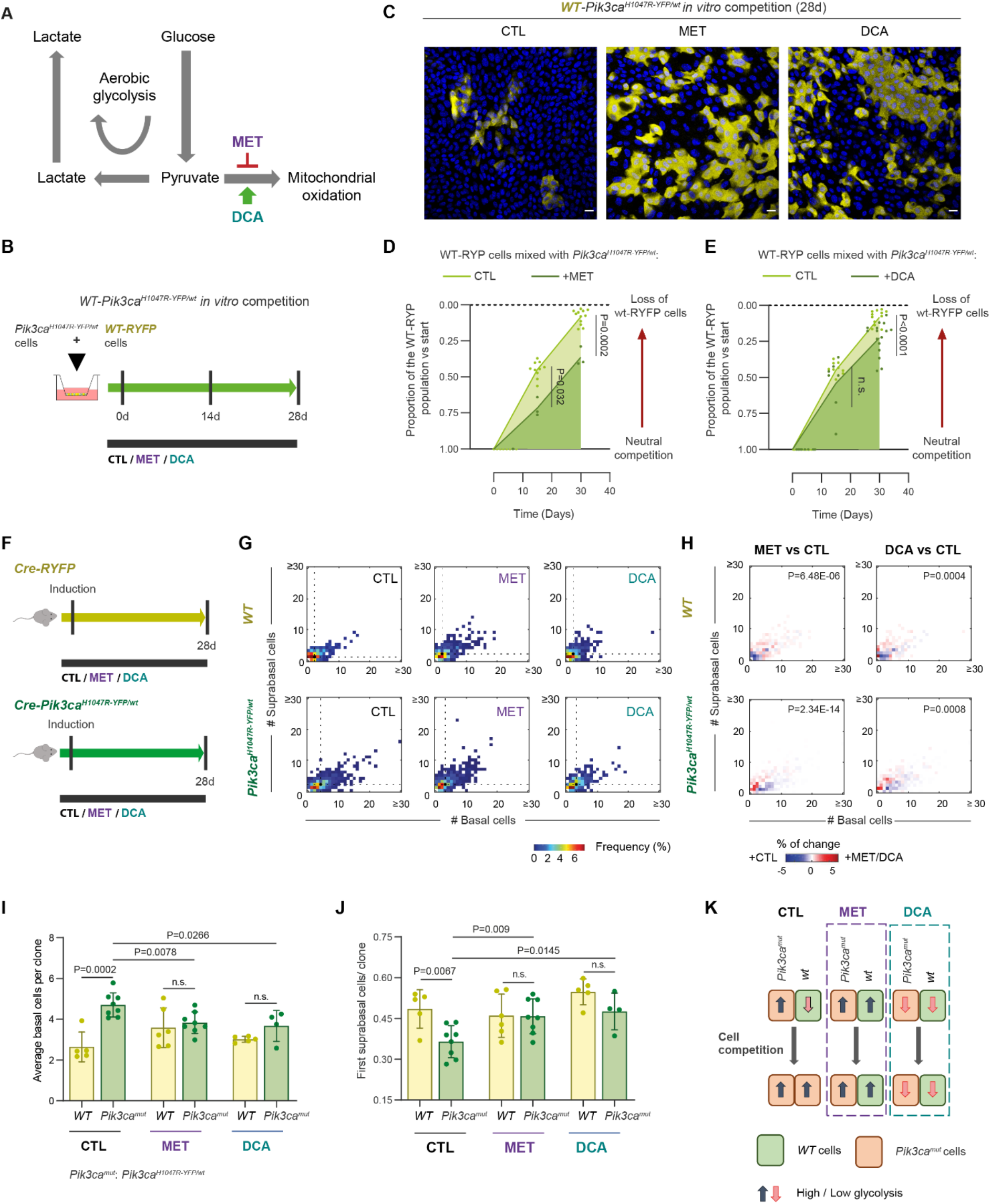
Metformin (MET) and DCA reduce *PIK3CA^H1047R-YFP/wt^* advantage *in vitro* and *in vivo*. (A) Scheme of energy metabolism with MET inhibiting mitochondrial oxidation (red arrow) and DCA activating glucose oxidation (green arrow). (B) *In vitro* cell competition assay. *In vitro* induced *Rosa26^flYFP/flYFP^* primary esophageal keratinocytes (*WT-RYFP*) were mixed with *in vitro* induced *PIK3CA^H1047R-YFP/wt^* cells or uninduced controls from the same animals. Once a confluent culture is achieved, cells were treated for 28 days in culture with minimal FAD medium +/− MET 2.5 mM or +/− DCA 25 mM. Samples were collected at the start of the treatment and at 28 days and the proportion of *WT-RYFP* at the end of the experiment *versus* at the beginning was calculated. (C) Confocal z stack image representative of the co-culture of *WT-RYFP* cells and *PIK3CA^H1047R-YFP/wt^* after 28 days of continuous treatment in minimal FAD medium or minimal FAD medium with MET 2.5 mM or DCA 25 mM. An optical section through the basal cell layer is shown. YFP immunofluorescence (yellow), nuclei are stained with DAPI (blue). Scale bar, 20 μm. (D-E) *In vitro* cell competition assays. Proportion of *WT-RYFP* cells, mixed either with induced *PIK3CA^H1047R-YFP/wt^* cells or uninduced controls, *versus* the start of the experiment, at the specified time points. Cells were treated in minimal FAD medium or minimal FAD medium with MET 2.5 mM (B) or DCA 25 mM (C) for the duration of the experiment. Each dot represents a primary culture from an animal. n.s., not significant. n=3-16 primary cultures coming from different animals. Two-tailed ratio paired *t*-test. (F) Experimental protocol. *Cre-RYFP* reporter mice and *Cre-PIK3CA^H1047R-YFP/wt^* mice were induced with β-naphthoflavone and tamoxifen. They were treated with MET or DCA for the duration of the experiment and collected 28 days after induction. (G) Heatmaps showing the frequency of clone sizes with the number of basal and first suprabasal cells indicated observed in animals from (F). Black dots and dashed lines indicate geometric median clone size. n=311-917 clones from 5-10 animals per condition (see Table S1 for numbers). (H) Heatmaps showing the differences between each treatment and control in *Cre-RYFP* (upper panels) or *Cre-PIK3CA^H1047R-YFP/wt^* (lower panels) animals. n=311-917 clones from 5-10 animals per condition (see Table S1 for numbers). 2D Kolmogorov-Smirnov test. (I) Average basal clone sizes for each strain and treatment from (F), considering all clones with at least one basal cell. Dots indicate the average clone size of a mouse. Average and standard deviation per condition are indicated. n=4-8 animals per condition (animals with more than 50 clones). n.s., not significant. Two-tailed unpaired *t*-test. (J) Average proportion of suprabasal cells per clone for each strain and treatment from (F), counting basal and first suprabasal cells. Each dot corresponds to one animal. Average and standard deviation per condition are indicated. n=4-8 animals per condition (animals with more than 50 clones). n.s., not significant. Two-tailed unpaired *t*-test. (K) Model showing the results relating cell competition and glycolysis. *PIK3CA^H1047R-YFP/wt^* cells have increased glycolysis and an advantage over wild type cells in a control situation. Activating glycolysis with MET or reducing glycolysis with DCA; both in wild type and mutant cells, reduces the competitive advantage of *PIK3CA^H1047R-YFP/wt^* cell.

Finally, we determined whether MET and DCA can both inhibit clonal expansion *in vivo*, using *Cre-Pik3ca^H1047R-YFP/wt^* and *Cre-RYFP* mice. Animals were induced and treated with MET or DCA for one month, when clone sizes were analyzed (**Figure 6F**). Both MET and DCA as separate treatments reduced mutant clone size *in vivo* and increased the proportion of differentiated cells towards wild type levels (**Figures 6G-J and S5B**). These results support the hypothesis that reducing glycolytic differences between *Pik3ca^H1047R-YFP/wt^* mutant and neighboring wild type cells, lowers the bias towards proliferation and hence the competitive fitness of mutant progenitors (**Figure 6K**).

**Supplementary Figure 5.**
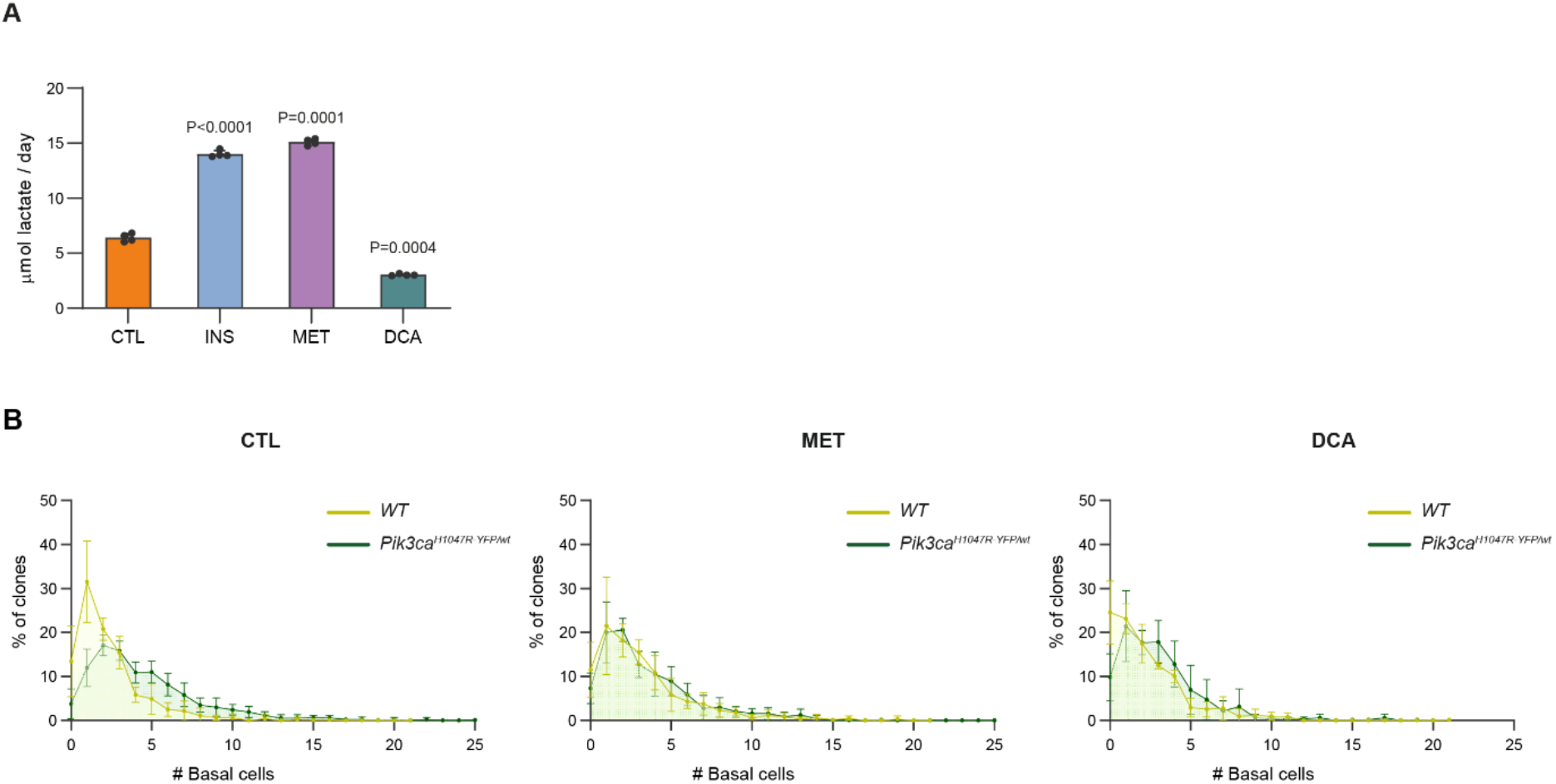
Related to Figure 6. Effect of treatments in lactate secretion *in vitro* and effect of MET and DCA treatments on basal cell distributions *in vivo*. (A) Lactate secretion in wild type primary esophageal keratinocytes treated in minimal FAD medium or minimal FAD medium with 5 μg/ml insulin (INS), 2.5 mM metformin (MET) or 25 mM DCA (DCA). (B) Distribution of basal cells per clone from *Cre-RYFP* reporter mice and *Cre-PIK3CA^H1047R-YFP/wt^* mice after 28 days of induction and treatment. n=311-917 clones from 5-10 animals per condition (see Table S1 for numbers).

## Discussion

The results presented here show that a subtle activation of the PI3K pathway caused by a heterozygous activating missense mutation in *Pik3a* is sufficient to drive clonal expansion in normal esophageal epithelium.

The cellular mechanism underpinning the competitive advantage of mutant *Pik3a* progenitors is a small increase in the probability of generating mutant progenitors over differentiated daughters per division, with no detectable acceleration in the cell cycle. A similar change in mutant progenitor dynamics, an increase in the proportion of proliferating *versus* differentiating cells per average cell division, occurs with a Notch inhibiting mutant and mutant *Trp53* in the mouse esophagus and skin respectively (Alcolea et al., 2014; Murai et al., 2018). It is striking that three disparate mutations under positive selection in human esophagus all result in a similar alteration in mutant cell dynamics. These observations indicate that altering progenitor cell fate is the common mechanism hijacked by mutations in different pathways to expand in squamous epithelia. Once clones collide with others of similar fitness progenitor cell fate reverts towards balanced production of progenitor and differentiating cells (Colom et al., 2020). A small fate imbalance towards proliferating cells also occurs in high grade dysplasias and carcinomas in the mouse esophagus (Frede et al., 2016). However, while esophageal tumors have the potential to grow, competition with other mutant clones in non-transformed tissue is constrained by the limited space available for mutant clone expansion (Colom et al., 2020; Frede et al., 2016).

We confirmed that in primary esophageal keratinocytes, *Pik3ca^H1047R^* expression activates glycolysis, as previously described in cell lines (Hu et al., 2016; Ilic et al., 2017; Jiang et al., 2018). However, the molecular basis of the effect of *Pik3ca^H1047R-YFP/wt^* on esophageal progenitor cell fate remains to be elucidated. Several studies of keratinocytes in the epidermis of the skin, have observed a link between the level of glycolysis and regulation of proliferation *versus* differentiation. For example, differentiation is increased by downregulation of glycolysis by activation of the aryl hydrocarbon receptor or inhibition of enolase (Sutter et al., 2019). Similarly, deletion of the glucose transporter, *Glut1,* results in decreased proliferation and increased reactive oxygen species (ROS) (Zhang et al., 2018). Elevation of ROS drives keratinocyte differentiation in both skin and esophagus (Fernandez-Antoran et al., 2019; Hamanaka et al., 2013). Finally, HIF1α also promotes proliferation in human skin keratinocytes (Kim et al., 2018). Thus, our results suggest that the regulatory role for HIF1α and glycolysis in keratinocyte differentiation may be a common feature of stratified epithelia.

The expansion of *Pik3ca^H1047R-YFP/wt^* clones depends not only on the mutant cell phenotype but also the activity of the PI3K signaling axis and metabolic state of the adjacent wild type cells. Reducing the differences in PI3K/HIF1α/aerobic glycolysis axis between mutant and wild type cells, attenuates the competitive advantage of the mutant (**Figure 7**). These observations parallel those of mutations that activate PI3K signaling in *Drosophila*, which confer a fitness advantage that may be modulated by alterations in insulin exposure or metformin (Nowak et al., 2013; Sanaki et al., 2020). Nevertheless, the relationship between aerobic glycolysis and competitive fitness seems context-dependent, resulting in elimination of glycolytic mutant cells in the mouse intestine but conferring a fitness advantage in the *Drosophila* wing disc (Banreti and Meier, 2020; Kon et al., 2017). Likewise, strong activation of PI3K signaling by biallelic expression of *Pik3ca^H1047R/H1047R^* in keratinocytes in the epidermis leads to their elimination by differentiation (Ying et al., 2018).

**Figure 7.**
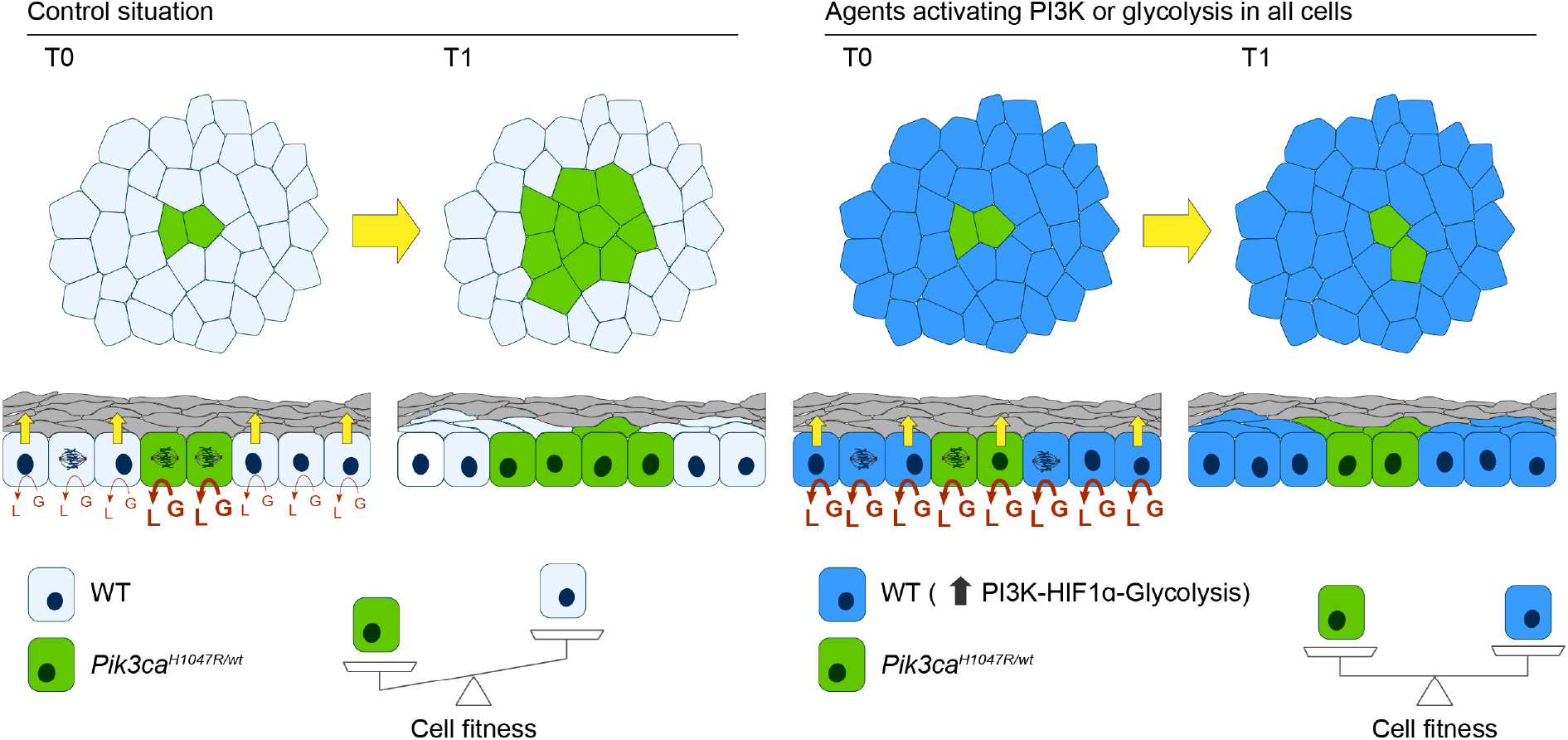
Model for the modulation of the competitive advantage *PIK3CA^H1047R-YFP/wt^* cells by interventions activating the PI3K pathway or glycolysis. Upon mutation, *PIK3CA^H1047R-YFP/wt^* cells have increased PI3K/HIF1α/Glycolysis pathway activity over their wild type neighbors and expand in the tissue due to a reduction in cell differentiation. Interventions activating the PI3K/HIF1α/Glycolysis pathway in wild type cells or inhibiting it in mutant cells, level the cell competition and reduce *PIK3CA^H1047R-YFP/wt^* mutant cell expansion. L: Lactate and G: Glucose.

Our findings hint that metabolic disease states such as insulin deficiency in type 1 diabetes or treatments such as long-term metformin administration, may alter competitive selection of signaling mutants in normal tissues, by modulating the advantage of clones with mutations that activate the PI3K pathway. Such remodeling of the normal tissue landscape may impact on the risk of neoplasia and represent a potential point of intervention in cancer prevention (Bradley et al., 2018; Carstensen et al., 2016; Yu et al., 2019). Beyond cancer, it is possible that part of the aging phenotype is due to the colonization of normal tissues by mutant clones (Blokzijl et al., 2016; Martincorena et al., 2018). If this is so, Metformin, currently in clinical trials as an ‘anti-aging’ drug, may have unexpected benefits in suppressing the expansion of a subset of mutant cell clones in normal adult epithelia (Barzilai et al., 2016).

## Experimental Model and Subject Details

### *Pik3ca^fl-H1047R-T2A-EYFP-NLS/wt^* generation

*Pik3ca^fl-H1047R-T2A-EYFP-NLS/wt^* knock-in mice were generated by Taconic Biosciences (Hudson, NY). In the targeting vector, exon 20 (including the splice acceptor site of intron 19), the endogenous STOP sequence and the 3'UTR region of the wild type *Pik3ca* gene were flanked by *loxP* sites. A second *Pik3ca* exon 20 including the *Pik3ca^H1047R^* mutation was introduced 3’ of the distal *loxP* site. Between the last amino acid and the translation termination codon of the duplicated exon 20 the following sequences were inserted: a self-cleaving T2A peptide, an EYFP fused to a Nuclear Localization Signal (NLS), a STOP cassette and the 3′UTR region of the *Pik3ca* gene. Two positive selection markers were also introduced. A Neomycin resistance gene flanked by Frt sites was inserted 3′ of the 5′ *loxP* site before exon 20. A Puromycin resistance gene flanked by F3 sites was placed downstream of the 3′UTR from the duplicated region. An additional polyadenylation signal (hGHpA: human Growth Hormone polyadenylation signal) was inserted between the 3’ UTR and the distal *loxP* site in order to prevent downstream transcription of the mutated *Pik3ca^H1047R^* exon 20 before Cre recombination. Finally, the vector included a distal thymidine kinase (Tk) gene at 3’ end for negative selection. The targeting vector was generated using BAC clones from the C57BL/6J RPCIB-731 BAC library and transfected into the Taconic Biosciences C57BL/6N Tac ES cell line. Homologous recombinant clones were isolated using double positive (NeoR and PuroR) and negative (Tk) selections. The appropriate insertion of the vector sequence was assessed by PCR. The conditional knock-in allele was obtained after *in vivo* Flp-mediated removal of the selection markers. This allele expresses the wild type p110α protein. The presence of the hGHpA cassette downstream of the wild type exon 20 prevents transcription of the mutated H1047R exon 20 and the EYFP before Cre recombination. The constitutive knock-in allele is obtained after Cre-mediated deletion of wild type exon 20 and the hGHpA. This allele expresses a chimeric transcript harboring the mutated p110α^H1047R^ protein fused to the T2A sequence and the EYFP open reading frame including the NLS. The expected co-translational cleavage at the T2A sequences result in co-expression of the mutated p110α^H1047R^ and EYFP proteins under the control of the endogenous *Pik3ca* promoter.

### Mouse experiments

All experiments were approved by the local ethical review committees at the Wellcome Sanger Institute, and conducted according to Home Office project licenses PPL70/7543, P14FED054 and PF4639B40. Animals were maintained on a C57/Bl6 genetic background, housed in individually ventilated cages and fed on standard chow. Experiments were carried out with male and female animals and no gender specific differences were observed. For lineage tracing experiments, the relevant floxed reporter lines were crossed onto the *Ahcre^ERT^* strain in which transcription from a transgenic CYP1A1 (arylhydrocarbon receptor, Ah) promoter is normally tightly repressed until is activated by β-napthoflavone (Kemp et al., 2004). In this model tamoxifen promotes the nuclear translocation of Cre^ERT^ protein to mediate recombination. For lineage tracing of control clones, the *Rosa26^flYFP/wt^* mice which express yellow fluorescent protein (YFP) from the constitutively active Rosa 26 locus were used (Srinivas et al., 2001). To assess the mutant and wild type clone growth in the same mouse, *Rosa26^flConfetti^* animals were used (Snippert et al., 2010). Homozygous *Ahcre^ERT^Rosa26^flConfetti^* animals were crossed onto *Pik3ca^fl-H1047R-T2A-EYFP-NLS/fl-H1047R-T2A-EYFP-NLS^* to generate *Ahcre^ERT^Rosa26^flConfetti^-Pik3ca^fl-H1047R-T2A-EYFP-NLS/wt^*. Following induction this strain yields cells expressing just one of 4 colors of reporter from the *Rosa26* locus (wild type clones), just the YFP reporter of Pik3ca mutant cells (*Pik3ca^H1047R-YFP/wt^* clones) or both (*Pik3ca^H1047R-YFP/wt^*-Confetti clones) (**Figure S2B**). Because GFP, YFP and CFP clones are recognized by the same anti-GFP antibody used to detect the EYFP co-expressed with the mutation, only RFP expressing clones (wild type or mutant) were analyzed. Where indicated, mice were treated with a well-tolerated dose of metformin (MET, Sigma-Aldrich 317240, 2 mg/ml) or sodium dichloroacetate (DCA, Sigma-Aldrich 347795, 0.5 mg/ml) in drinking water containing 15% Ribena to enhance palatability, for the duration of the experiment. In those cases, the results were compared to control mice treated with 15% Ribena in water. For the analysis of *Pik3ca^H1047R-YFP/wt^* mutant clones in a hypoinsulinemic background, *Ahcre^ERT^Pik3ca^fl-H1047R-T2A-EYFP-NLS/fl-H1047R-T2A-EYFP-NLS^* female mice were crossed with heterozygous *C57BL/6-Ins2^Akita^/J* males (*Akita^Het^*) (The Jackson Laboratory US) in order to produce *Ahcre^ERT^-Pik3ca^fl-H1047R-T2A- EYFP-NLS/wt^-Ins2^Akita/wt^* and *AhCre^ERT^-Pik3ca^fl-H1047R-T2A-EYFP-NLS/wt^-Ins2^wt/wt^* littermates. The development of diabetes phenotype was confirmed by analyzing the presence of glucose in urine, a hallmark of diabetes, at the start and end of the experiments (**Figure S3D**) using SURERS2/50 strips (SureScreen Diagnostics, UK) according to the manufacturer’s instructions.

### Lineage tracing

Low frequency expression of EYFP in the mouse esophagus was achieved by inducing transgenic animals aged 10–16 weeks with a single intraperitoneal (i.p.) of 80 mg/kg β-naphthoflavone and 1 mg tamoxifen and 80 mg/kg β-naphthoflavone (*Ahcre^ERT^Rosa26^flConfetti^-Pik3ca^fl-H1047R-T2A-EYFP-NLS/wt^*) and 0.25 mg tamoxifen (rest of mouse strains). Following induction, between three and eight mice per time point were culled and the esophagus collected. Time points analyzed include 10 days, 1, 3 and 6 months after the induction. As the expression from the endogenous *Pik3ca* locus is very low, immunofluorescence was necessary in order to detect the RYFP expression (**Figure 2D**). The total number of clones quantified for each figure can be found in **Table S1**. Normalized, clone-size distributions were built for each experimental condition and time point from the observed relative frequencies *f_m,n_* of clones of a certain size, containing *m* basal and *n* suprabasal cells, resulting in two-dimensional histograms, (displayed as heatmaps using CloneSizeFreq_2Dheat package (https://github.com/gp10/CloneSizeFreq_2Dheat). A 2D histogram of the residuals or differences observed between conditions in the relative frequencies of each particular clone size (i.e., each cell on the grid) was generated when appropriate.

**Table S1.** Number of quantified clones per condition and time-point.

| Fig 3 | *R26<sup>YFP/wt</sup>* |  | *Pik3ca<sup>H1047R/wt</sup>* |  |
| --- | --- | --- | --- | --- |
|  | Clones quantified | Animals analysed | Clones quantified | Animals analysed |
| 10 days | 1522 | 6 | 659 | 4 |
| 28 days | 710 | 5 | 659 | 6 |
| 84 days | 457 | 5 | 629 | 6 |
| 168 days | 317 | 6 | 563 | 5 |

| Fig S2 | *R26<sup>RFP/wt</sup> Pik3ca<sup>wt/wt</sup>* |  | *R26<sup>RFP/wt</sup> Pik3ca<sup>H1047R/wt</sup>* |  |
| --- | --- | --- | --- | --- |
|  | Clones quantified | Animals analysed | Clones quantified | Animals analysed |
| 10 days | 316 | 3 | 153 | 3 |
| 28 days | 212 | 2 | 101 | 2 |
| 84 days | 204 | 2 | 51 | 2 |

| Fig 4 | *Pik3ca<sup>H1047R/wt</sup>* |  |
| --- | --- | --- |
| 28 days | Clones quantified | Animals analysed |
| CTL | 402 | 4 |
| AKITA | 257 | 4 |

| Fig 6 | *R26<sup>YFP/wt</sup>* |  | *Pik3ca<sup>H1047R/wt</sup>* |  |
| --- | --- | --- | --- | --- |
|  | Clones quantified | Animals analysed | Clones quantified | Animals analysed |
| 28 days |  |  |  |  |
| CTL | 431 | 5 | 799 | 8 |
| MET | 597 | 6 | 917 | 10 |
| DCA | 407 | 5 | 311 | 5 |

### Quantitative analysis and mathematical modelling

For details of quantitative analysis of wild type and *Pik3ca* mutant progenitor cell lineage tracing data and the dynamics of mutant cells in the suprabasal cell layers, see Supplementary Text. Code used for this analysis has been made publicly available and can be found at https://github.com/gp10/DriverClonALTfate.

### EdU *in vivo* proliferating cell analysis

One month after lineage tracing induction, 10 μg of EdU in PBS was administered by intraperitoneal injection 1 h before culling. Tissues were collected and stained with EdU-Click-iT kit and immunofluorescence as explained below. EdU-positive basal cells were quantified from a minimum of 10 z-stack images.

### Primary keratinocyte 3D culture

After removing the muscle layer with fine forceps, esophageal explants were placed onto a transparent ThinCert™ insert (Greiner Bio-One) with the epithelium facing upward and the submucosa stretched over the membrane, and cultured in complete FAD medium (50:50) 4.5 g/L D-Glucose, Pyruvate, L-Glutamine D-MEM (Invitrogen 11971-025): D-MEM/F12 (Invitrogen 31330-038), supplemented with 5 μg/ml insulin (Sigma-Aldrich I5500), 1.8 × 10^−4^ M adenine (Sigma-Aldrich A3159), 0.5 μg/ml hydrocortisone (Calbiochem 386698), 1 ×10^−10^ M cholera toxin (Sigma-Aldrich C8052), 10 ng/ml Epidermal Growth Factor (EGF, PeproTech EC Ltd 100-15), 5% fetal calf serum (PAA Laboratories A15-041), 5% Penicillin-Streptomycin (Sigma Aldrich, P0781) and 5 μg/ml Apo-Transferrin (Sigma-Aldrich T2036). Explants were removed after 7 days once keratinocytes have covered half of the membrane. Media was changed every three days. Minimal FAD medium without Cholera toxin, epidermal growth factor, insulin and hydrocortisone was used from two weeks after stablishing the culture.

### Adenoviral infection

To establish mouse primary keratinocyte 3D cultures from *Pik3ca^fl-H1047R-T2A-EYFP-NLS/wt^* mice and *Rosa26^flYFP/flYFP^*, cells were infected with *Cre*-expressing adenovirus (Ad-CMV-iCre, Vectorbiolabs, #1045 US) or Null adenovirus (Ad-CMV-Null, Vectorbiolabs, #1300 US). Briefly, cells were incubated with adenovirus-containing medium supplemented with Polybrene (Sigma Aldrich, # H9268) (4 μg/ml) for 24 h at 37°C, 5% CO_2_. Cells were washed and fresh medium was added. Infection rates were > 90%.

### Cell competition assays

Fully induced Rosa26^flYFP/flYFP^ cultures (*WT-RYFP*) were trypsinized and mixed, 1:1 for FACS analysis or 1:3 for microscopy, with *Pik3ca^fl-H1047R-T2A-EYFP-NLS/wt^* either fully induced with Cre-expressing adenovirus or uninduced (infected with null adenovirus). After one week, when the cultures are fully confluent, cholera toxin, epidermal growth factor, insulin and hydrocortisone were removed from the medium and starting time point was collected and analyzed by flow cytometry. At the time points specified, cells were collected and analyzed by flow cytometry or fixed for microscopy. Where indicated, mixed cultures were treated with 5 μg/ml insulin (Sigma-Aldrich I5500), 10 μM PX478 (Cambridge Bioscience 10005189), 2.5 mM metformin (Sigma-Aldrich 317240) or 25 mM sodium dichloroacetate (Sigma-Aldrich 347795).

### Flow cytometry

Keratinocyte cultures were detached by incubation with 0.05% Trypsin-EDTA for 20 min at 37°C 5% CO_2_. Cells were pelleted for 5 min at 650 g and resuspended in PBS to be immediately analyzed using a BD LSRFortessa. Were suprabasal and basal cells needed to be quantified, 2% PFA fixed cells were incubated in blocking solution (0.1% BSA 0.5 mM EDTA in PBS) for 15 min and then with anti-ITGA6-647 antibody (1:125, Biolegend, UK 313610) in blocking solution for 45 min at RT. YFP fluorescence was collected using the 488 nm laser and the 530/30 bandpass filter and ITGA6-647 fluorescence was collected using the 640 nm laser and the 670/14 bandpass filter. Data was analyzed using FlowJo software (version 10.5.3). Basal cells were defined as ITGA6 positive cells and suprabasal cells as ITGA6 negative cells.

### Immunofluorescence and microscopy

For wholemount staining, the mouse esophagus was opened longitudinally, the muscle layer was removed and the epithelium was incubated for 1 h and 30 min in 20 mM EDTA-PBS at 37°C. The epithelium was peeled from underlying tissue and fixed in 4% paraformaldehyde in PBS for 30 min. Wholemounts were blocked for 1 h in blocking buffer (0.5% bovine serum albumin, 0.25% fish skin gelatin, 1% Triton X-100 and 10% donkey serum) in PHEM buffer (60 mM PIPES, 25 mM HEPES, 10 mM EGTA, and 4 mM MgSO_4_·7H_2_O). Anti-GFP antibody (1:4000, Life technologies A10262) was incubated over 3-5 days using blocking buffer, followed by several washes over 24 h with 0.2% Tween-20 in PHEM buffer. Where indicated, an additional overnight incubation with anti-caspase-3 (1:500, Abcam ab2302) was performed. Finally, samples were incubated for 24 h with 1 μg/ml DAPI and secondary antibody anti-chicken (1:2000, Jackson ImmunoResearch 703-545-155) in blocking buffer. Clones were imaged on an SP8 Leica confocal microscope. Whole tissue or large tissue area images were obtained in most cases with the 20x objective with 1× digital zoom, optimal pinhole and line average, speed 600 hz and a pixel size of 0.5678μm/pixel. In YFP tissues at long time points, clones were manually detected in the microscope and individually imaged using 40× objective with 0.75× digital zoom, optimal pinhole and line average, speed 600 hz and a pixel size of 0.3788 μm/pixel. The numbers of basal and suprabasal cells in each clone were counted manually. Representative images were produced by selecting a 120×120 μm area in the images obtained as previously stated. Images were processed using ImageJ software adjusting brightness and contrast and applying a Gaussian blur of 1. EdU incorporation was detected with Click-iT chemistry kit according to the manufacturer’s instructions (Invitrogen) using 555 Alexa Fluor azides. Confocal images for EdU-GFP staining were acquired on a Leica TCS SP8 confocal microscope (objective 20×; optimal pinhole and line average; speed 600 Hz; resolution 1024 × 1024, zoom ×2). Images were processed using ImageJ software adjusting brightness and contrast and applying a Gaussian blur of 1. For *in vitro* culture staining, inserts were fixed in 4% paraformaldehyde in PBS for 30 min, then blocked for 30min in blocking buffer and incubated overnight with anti-GFP antibody (1:1000, Life technologies A10262) in blocking buffer followed by 4x 15 min washes with 0.2% Tween-20 in PHEM buffer. Finally, samples were incubated for 2h with 1 μg/ml DAPI and secondary antibody anti-chicken (1:500, Jackson ImmunoResearch 703-545-155) in blocking buffer. Afterwards, inserts were washed 4× 15 min with 0.2% Tween-20 in PHEM buffer and mounted in Vectashield (Vector Laboratories). Cultures were imaged on an SP8 Leica confocal microscope, obtained with the 40× objective with 0.75× digital zoom, optimal pinhole and line average, speed 600 hz and a pixel size of 0.1893 μm/pixel.

### RNA isolation and RNA sequencing

Total RNA was extracted from 3D cultures of mouse primary keratinocytes after 1 week in FAD medium supplemented with fetal calf serum, apo-transferrin and Penicillin/Streptomycin, with or without insulin. RNA was extracted using RNeasy Micro Kit (QIAGEN, UK), following the manufacturer’s recommendations. Briefly, cells were washed with cold Hank’s Balanced Salt Solution-HBSS (GIBCO, UK) and then lysis buffer was added directly to the insert. The integrity of total RNA was determined by Qubit RNA Assay Kit (Invitrogen, UK). For RNA-seq, libraries were prepared in an automated fashion using an Agilent Bravo robot with a KAPA Standard mRNA-Seq Kit (KAPA BIOSYSTEMS). In house adaptors were ligated to 100-300 bp fragments of dsDNA. All the samples were then subjected to 10 PCR cycles using sanger_168 tag set of primers and paired-end sequencing was performed on Illumina HiSeq 2500 with 75 bp read length. Reads were mapped using STAR 2.5.3a, the alignment files were sorted and duplicate-marked using Biobambam2 2.0.54, and the read summarization performed by the htseq-count script from version 0.6.1p1 of the HTSeq framework (Anders et al., 2015; Dobin et al., 2013). For GSEA analysis raw counts were normalized by the median of ratios method (Love et al., 2014). Gene set enrichment was analyzed with GSEA software (Subramanian et al., 2005) using the Hallmarks gene sets of the Molecular Signature Database (MSigDB) version 4.0 provided by the Broad Institute (http://www.broad.mit.edu/gsea/), following the standard procedure described on the GSEA user guide (http://www.broadinstitute.org/gsea/doc/GSEAUserGuideFrame.html). The FDR for GSEA is the estimated probability that a gene set with a given NES (normalized enrichment score) represents a false-positive finding, and an FDR<0.25 is considered to be statistically significant for GSEA. Differential gene expression was analyzed using the DEBrowser tool (https://debrowser.umassmed.edu/) with which we performed a DESeq2 analysis (Love et al., 2014) filtering the low counts to remove genes with less than 2 cpm in at least 2 samples. Parametric’ fitting of dispersions to the mean intensity was used with the likelihood ratio test on the difference in deviance between a full and reduced model formula (defined by nbinomLRT). An adjusted p-value cut-off of 0.05 were used to select significantly different expressed genes. Heatmaps were generated from the TPM values and build using ClustVis (https://biit.cs.ut.ee/clustvis/) and Morpheus tools (https://software.broadinstitute.org/morpheus/), significance was calculated from the adjusted p-value obtained in the DE analysis. Kyoto Encyclopedia of Genes and Genomes (KEGG) pathway enrichment analysis was performed uploading the significantly upregulated gene list (p<0.05) into the Enrichr tool (https://amp.pharm.mssm.edu/Enrichr/) (Kuleshov et al., 2016). Venn diagrams were generated using the Venn Diagrams tool (https://www.biotools.fr/misc/venny). MA plots were generated using GraphPad Prism 8.

### Validation of *Pik3ca^H1047R^* mutant construct in NIH3T3

DNA that reflects the recombined *Pik3ca^H1047R-YFP^* allele (*PIK3CA*^H1047R^ fused to self-cleaving peptide P2A and GFP) was chemically synthesized (GenScript USA Inc.). cDNA encoding murine p110α^wild^ ^type^, p110α^H1047R^ and p110α^H1047R^-P2A-GFP were amplified by PCR using IMAGE clone (Image ID 40141870/IRCL34 C10 (M13R), Source Bioscience) for wild type p110α or above cDNA for mutants and subcloned into the pCS2+ expression vector. NIH3T3 cells were transiently transfected with these constructs using Lipofectamine 2000 (Thermo Fisher 11668030) according to manufacturer’s instruction. Cells were serum-starved by culture in DMEM containing 0.3% serum for 23 h. Serum-starved NIH3T3 cells were lysed in buffer containing 20 mM Hepes NaOH pH 7.9, 10% Glycerol, 0.4 M NaCl, 0.5% NP-40, 0.2 mM EDTA, 0.01% halt protease and phosphatase inhibitor (ThermoFisher Scientific, cat #78415). Protein concentrations were measured using standard Bradford protein assays (BioRAD QuickSTART™ Bradford Dye Reagents, cat.no.500-0202). Lysates were mixed with equal amount of 2× loading buffer (100 mM Tris-HCl pH 6.8, 4% SDS, 20% Glycerol, Bromophenol blue and 0.2% β-mercaptoethanol) and boiled at 96°C for 5 min. Samples were loaded onto a 7.5 or 10% of SDS-polyacrylamide gel. Proteins were separated by electrophoresis and transferred onto Immobilon-P membrane (pore size 0.45 μm, Millipore IPVH00010). Membranes were incubated in blocking buffer (5% dried skimmed milk, PBS, 0.1% Tween-20) at room temperature for 1 h and then with primary antibodies diluted in blocking buffer for 1 h at room temperature or overnight at 4°C on a rocking platform. After washing in PBST (0.1% Tween-20, PBS) three times, HRP conjugated secondary antibodies diluted in 0.5% skimmed milk in PBST were applied to the membrane for 10 min at room temperature on a rocking platform followed by four washes in PBST 20 min each. Washing and secondary antibodies steps were performed using SNAP id protein detection system (Sigma-Aldrich). Proteins were detected using Immobilon Western Chemiluminescent HRP substrate (Millipore WBLUC0500) or ECL blotting reagents (GE Healthcare GERPN2109).

### Immune capillary electrophoresis

For protein phosphorylation analysis, 3D cultures were starved in FAD medium 0.1% FCS without cholera toxin, epidermal growth factor, insulin and hydrocortisone for 16 h at 37°C 5% CO_2_. Then treated for 15 min in the same starving medium or FAD medium with cholera toxin, epidermal growth factor, insulin, hydrocortisone and 20% FCS with or without LY294002 (50 μM). Cultures were lysed in ice-cold RIPA buffer (Thermo Scientific, UK) containing protease and phosphatase inhibitors. Plates were frozen at −80°C and thawed on ice, scraped and passed twice through a Qiashredder (2 min centrifuged at maximum speed), then incubated 1 h on ice vortexing every 15 min. Then lysates were centrifuged at 14000 g for 20 min at 4°C. The supernatant was collected for analysis. Total protein quantification was performed using Pierce BCA Protein Assay Kit (Thermo Scientific, UK). Immune capillary electrophoresis was performed using Wes Simple™ (ProteinSimple, USA) following manufacturer’s instructions. Primary antibodies used were p-AktS493 (New England Biolabs 4060S, 1:150), Akt (New England Biolabs 4691S, 1:100), p-PRAS40 (New England Biolabs 2691T, 1:300) and PRAS40 (New England Biolabs 2997T, 1:150).

### Respirometry experiments

Fully recombined *Pik3ca^H1047R/wt^* or uninduced (adenovirus-null infected) parallel cultures from the same mice were treated for at least 1 week in FAD medium supplemented with fetal calf serum, apo-transferrin and Penicillin/Streptomycin, with or without insulin. At the moment of the experiment, 5 mm circular sections of each culture were obtained using a biopsy punch (Stiefel BC-BI-1600) and transferred to XFe24 Cell Culture microplate wells with the basal layer facing up. 675 μl of bicarbonate-free DMEM (Sigma-Aldrich, D5030) supplemented with 25 mM glucose, 1 mM pyruvate, 4 mM glutamine, 5 μg/ml Apo-Transferrin with or without 5 μg/ml insulin (Sigma-Aldrich I5500) was added to each sample. To eliminate residual carbonic acid from medium, cells were incubated for at least 30 min at 37°C with atmospheric CO_2_ in a non-humidified incubator. OCR and ECAR were assayed in a Seahorse XF-24 extracellular flux analyzer by three measurement cycles of 2-min mix, 2-min wait, and 4-min measure. Then OCR/ECAR ratio was calculated. Five technical replicates and four biological replicates were collected per condition.

### Lactate measurement

Cultures were incubated for 3 days in the specified treatments. Medium was collected and lactate was analyzed using a commercial kit (DF16, Siemens Healthcare). All sample measurements were performed by the MRC MDU Mouse Biochemistry Laboratory.

### Quantification and Statistical Analysis

Unless otherwise specified, all data are expressed as mean values ± standard deviation. Differences between groups were assessed by 2-tailed unpaired *t*-test for normally distributed data or 2-tailed Mann-Whitney U test for skewed data, using GraphPad Prism software. For paired comparisons of clonal behavior across conditions, Peacock’s test was used, a two-dimensional extension of Kolmogorov-Smirnov test (implemented as kstest_2s_2d function for MATLAB). Exact p-values for statistical tests are specified in the figures. No statistical method was used to predetermine sample size. The experiments were not randomized. The investigators were not blinded to allocation during experiments and outcome assessment.

## Data Availability

Raw transcriptomic data can be viewed on https://www.ebi.ac.uk/ena using the accession numbers provided in STAR methods.

---

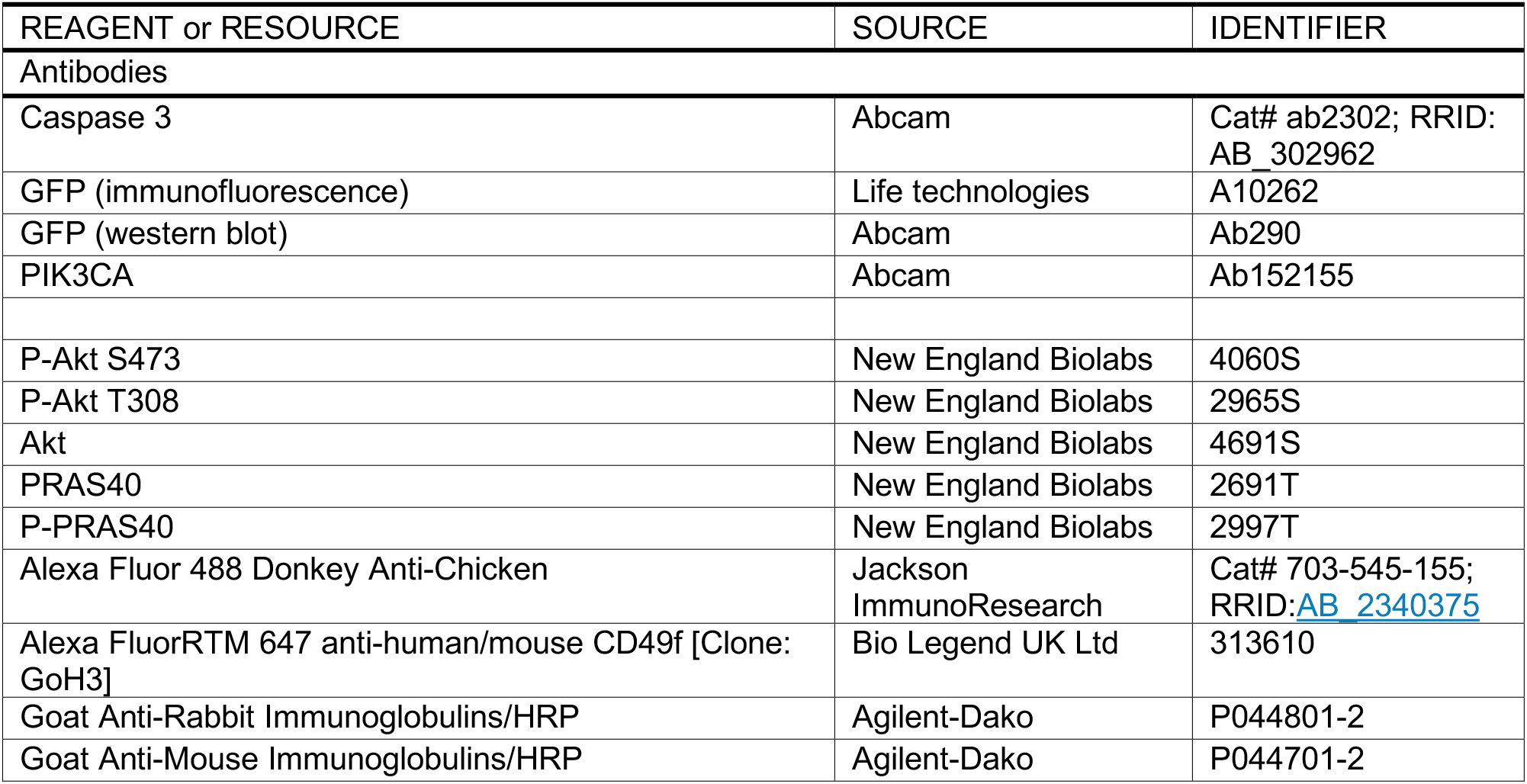
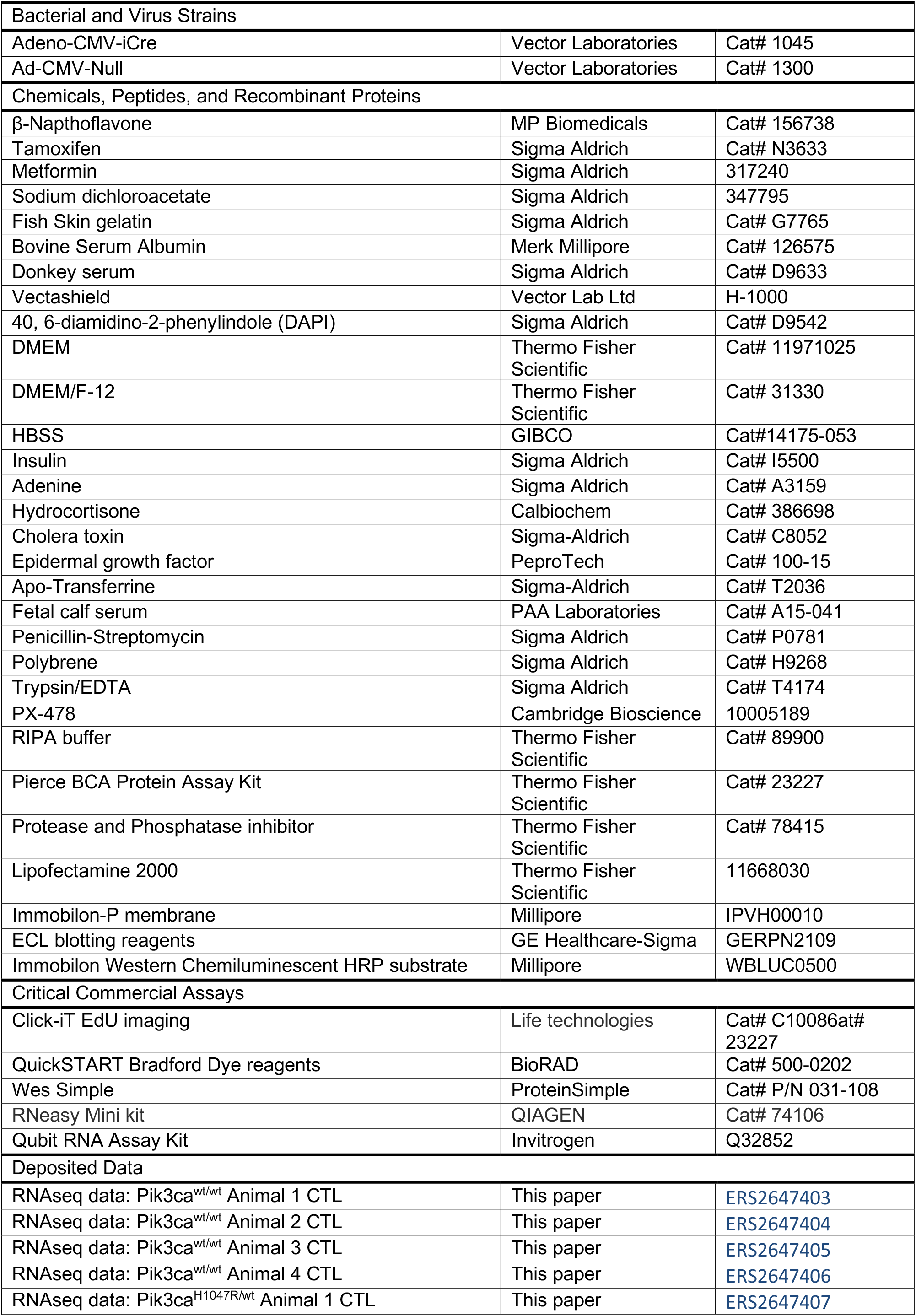
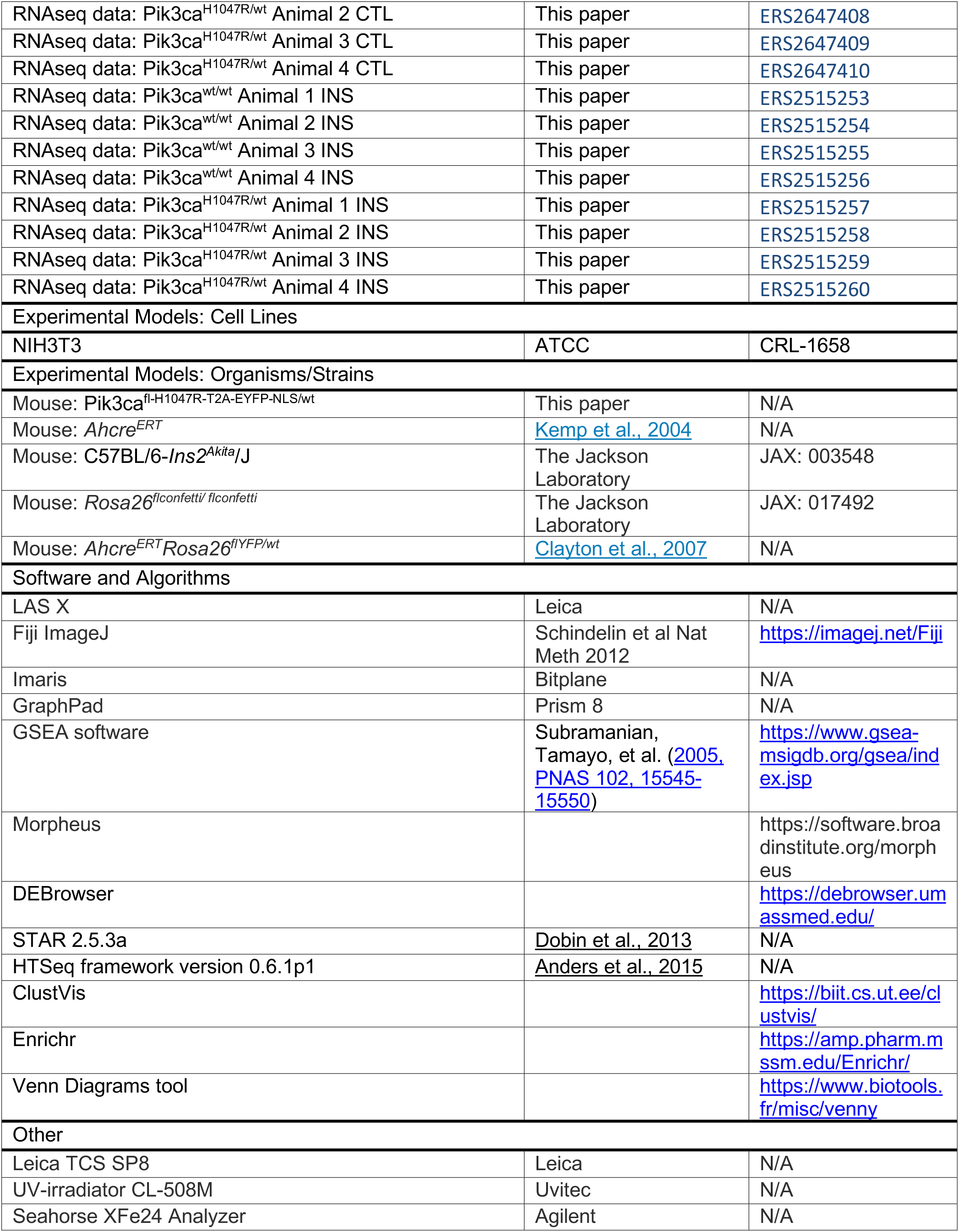
KEY RESOURCES TABLE.

## Acknowledgements

We thank Esther Choolun and Tom Metcalf for their assistance in *in vivo* experiments, and Wellcome Sanger Institute RSF facilities for technical support.

This work was supported by grants from the Wellcome Trust to the Wellcome Sanger Institute (098051 and 296194) and Cancer Research UK Programme Grants to P.H.J. (C609/A17257 and C609/A27326). A.H. benefited from the award of an EMBO long term fellowship. Work in the laboratory of B.V. is supported by Cancer Research UK (C23338/A25722) and the UK NIHR University College London Hospitals Biomedical Research Centre.

## Author Contributions

A.H. and B.C. performed the experiments with support from D.F-A. and C.B. K.M., B.V. and P.H.J. designed the mouse model and performed initial validations. G.P. analyzed clonal data. S.H.O. performed bioinformatics analysis. A.H., B.C. and P.H.J. wrote the manuscript with input from C.F. and B.V.

## Lead Contact and Materials Availability

Requests for reagent and resource sharing should be addressed to the Lead Contact, Philip H. Jones who will fulfil requests.

## Supplemental Information

<u>Document S1</u>. Figures S1–S6 and Table S1. (Embedded in this manuscript) <u>Table S2</u>. Source Data, Related to Figures 1, 3, 4, 5, 6a, and S1–S5.

<u>Document S2</u>. Supplementary Theory. (Embedded in this manuscript) <u>Supplementary video 1</u>.

2D simulation of the growth of wild type and *Pik3ca^H1047R/wt^* clones over 6 months starting from the same proportion of induced cells and following the parameters described in **Figure 3I**.

## SUPPLEMENTARY THEORY

This report provides a detailed description of the quantitative methods and modelling used to study mutant keratinocyte behavior in the murine esophagus. **Section 1** covers the analysis of wild type progenitor cell behavior. In **Section 2** we extend the methodology to test *Pik3ca* mutant progenitor cell dynamics. In **Section 3** we analyze cell dynamics in the suprabasal compartment for model validation.

### 1. Wild type progenitor cell dynamics

The murine esophageal epithelium consists of layers of keratinocytes maintained by a single type of proliferative cells (Doupe et al., 2012; Piedrafita et al., 2020). Progenitor (P-) cells reside in the deepest, basal layer where they divide regularly at a rate λ. The outcome of a given progenitor division is stochastic: with a certain probability it results in two proliferating daughter cells retaining the proliferative capacity (P+P) or two post-mitotic, differentiating cells (D+D), the remaining divisions yielding an asymmetric outcome (P+D). Upon differentiation, D-cells stratify (at rate Г) into the upper, suprabasal layers (transiting to S-cells), being ultimately shed into the lumen (at rate μ). This scenario is summarized by the single-progenitor (SP) model (**Figure 2B**):

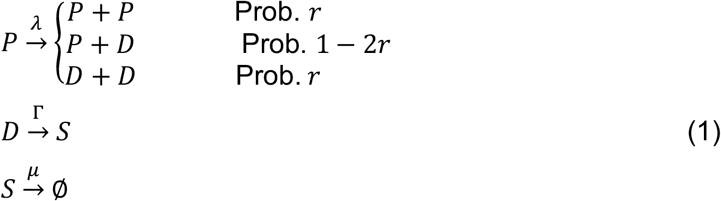

While the outcome of individual divisions is unpredictable, overall, the likelihood of the symmetric PP and DD division outcomes is balanced in adult wild type mice (setting the same probability, *r*). This ensures that on average half the progenitor cells go on to divide and half differentiate, so that the tissue remains homeostatic.

Under homeostasis, one can assume the proportion of proliferative basal cells, ρ, remains constant, and overall, the net rate at which cells are generated in the basal compartment is compensated by cell stratification and cell loss by shedding (**Figure 2A**). Then, the following relationships between the parameters can be established: *Г* = *λρ*/(1 − *ρ*), and *μ* = *λρ*(1 − *h*)/*h*, where *h* is the proportion of suprabasal cells relative to total cells. Alternatively, one can set *μ* = *λρ*/*m*, if we define *m* as the global ratio of suprabasal-to-basal cell populations.

Our lineage tracing data from *Cre-RYFP* (wild type) mice are consistent with key dynamical features of the SP model with balanced fates, in agreement with our more extensive work carried out previously using the same mouse strain and others (Clayton et al., 2007; Doupe et al., 2012; Piedrafita et al., 2020). First, persisting clones (i.e. those retaining at least one basal cell) show ever increasing sizes during the duration of the experiment. In particular, the average number of basal cells per persisting clone follows a linear growth over time (**Figure 3B**). Second, clones become increasingly heterogeneous in size, both in terms of the number of basal cells and total (basal + 1^st^ suprabasal) cells per clone, the distributions adopting a scaling behavior at late time points (**Figure 3D**). These are hallmarks of neutral clone competition, where some clones grow by chance at the expense of others that shrink, lose basal attachment and get ultimately extinct by shedding (**Figure 3H**).

In order to validate the dynamics of the wild type progenitor cells in our particular experimental setup, we proceeded to determine the values for the unknown SP-model parameters by fitting the experimental basal clone size distributions at the different time points (suprabasal cell numbers are not required for this since proliferative cells are confined to the basal compartment). For the division rate, we took as a prior the average value for and the distribution of cell-cycle time periods *t_cc_* inferred by H2B-GFP dilution chase experiments in (Piedrafita et al., 2020), i.e. < *λ* > = 2.9 week^−1^ and *t*_*cc*_~*τ*_*R*_ + Gam(*κ*, *ϑ*), where *τ*_R_ = 0.5 day^−1^ (refractory period) and Gam refers to a Gamma distribution with *κ* = 8 and *ϑ* = (1 − *τ*_R_ < *λ* >)/(*κ* < *λ* >). Notice that the election of this realistic description of the division rate will condition the modelling implementation as this cannot rely on Poisson processes that assume independent, underlying exponential events (see below). A maximum likelihood estimation (MLE) approach was then followed to infer the values of the two other remaining parameters, *r* and (which sets the value of Г in homeostasis), as explained below.

We performed a grid search spanning the range of all possible values for *r* and, and for every parameter set θ we estimated the log-likelihood value *l(θ ; x)* as in previous work (**Figure S2G**) (Murai et al., 2018; Piedrafita et al., 2020):

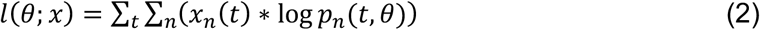

*x*_*n*_(*t*) is the observed frequency of clones with a certain basal size *n* at time *t*. In turn, *p*_*n*_(*t, θ*) refers to the probability of observing clones of that size at time *t* given the SP model with parameter values θ, and it was obtained by numerical solution (≥ 100,000 simulations) of the Master equation. In particular, in (Murai et al., 2018), this was computed by implementing a Markov-chain Monte Carlo method (Gillespie’s algorithm) (Gillespie, 1977; Gillespie, 1976). Here, we used an exact non-Markovian Monte Carlo analogue developed in (Piedrafita et al., 2020), which allows to account for the gamma-distributed cell cycle times. For convenience, both experimental and simulated clone sizes were binned in ranges increasing in powers of two given the large asymmetry of the distributions, so that in practice, *n* in the equation above stands for clones with a number of basal cells in the range (2^n-1^ + 1, 2^n^).

To discard possible biases due to the initial induction of post-mitotic cells (D- or S- cells), only clones with at least two basal cells were considered for the analysis. Also, a small fraction of clones at the latest time point (2 out of 317 clones in the wild type; 3 out of 563 clones in the mutant) were classified as outliers (i.e. having a number of basal cells ≫ 2,3 s.d. above the average of that time) and reassigned to the top size range *n* of non-outlier clones (Murai et al., 2018; Piedrafita et al., 2020). Finally, since there were large differences in our experimental sample size across time points (especially in the wild type: 1522, 710, 457 and 317 clones quantified at time 10 d, 28 d, 84 d and 168 d, respectively) and these can imprint uneven contributions to *l*(*θ* ; *x*) calculations (**Eq. 2**), here we followed a bootstrapping strategy: We computed *l*(*θ* ; *x*) repeatedly for different sample subsets containing a fixed number of clones *X* across time points, drawn by random permutation with replacement of the original sampled clones. In this way we ensured even weights from the different time points and more robust parameter estimates by averaging *l*(*θ* ; *x*) across subsamples.

Following this methodology, we obtained the following parameter estimates 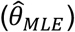 for the wild type progenitors (**Figure 3I**; **Figure S2G**):

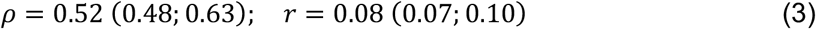

From these, we derive the stratification rate (recall homeostatic relationships above): Г = 3.14 (2.68; 4.94) week^−1^. In parentheses are 95% confidence interval bounds based on likelihood-ratio test (Pearson’s *χ*^2^ cutoff with 2 degrees of freedom). Altogether, these parameter values are in good agreement with those found previously from lineage tracing using inducible *Ah-Cre^ERT^* and *Lrig1-Cre^ERT^* based mice ({*ρ* = 0.56 (0.50; 0.89), *r* = 0.06 (0.04; 0.10)} and {*ρ* = 0.65 (0.50; 0.96), r = 0.10 (0.07; 0.15)}, respectively) (Piedrafita et al., 2020). It follows that using the SP model with the MLE values (**Eq. 3**) we obtained good fits on both the experimental average number of basal cells per clone (**Figure 3B**) and the distributions of basal clone sizes at the different time points (**Figure S2I**), corroborating the robust dynamics of wild type progenitors in the esophageal epithelium.

### 2. *Pik3ca^H1047R-YFP/wt^* progenitor cell dynamics

The dynamics of *Pik3ca^H1047R-YFP/wt^* mutant progenitor cells clearly differed from wild type cells as clones showed an accelerated growth over time in both the basal and suprabasal compartments (**Figure 3B-D**). The percentage of EdU^+^ basal cells among the induced *Pik3ca^H1047R-YFP/wt^* population was similar to wild type (**Figure 3F**) and comparable to measurements done before in mouse esophagus (Fernandez-Antoran et al., 2019). This argues against changes in the rate of mutant cell division *λ*. Yet, in principle dynamics might still be explained by a SP model with balanced fates if *Pik3ca^H1047R-YFP/wt^* progenitors experienced changes in some of the other parameters (e.g. *r* or Г). Alternatively, it could be that *Pik3ca^H1047R-YFP/wt^* clone behavior responds to an imbalance in mutant progenitor division outcomes that favors proliferating daughter cells over differentiating progeny (i.e. PP symmetric division outcome being more likely than DD) (**Figure 2B**). This later scenario has been shown to explain the phenotype of some other inducible mutants in squamous epithelium such as *DN-Maml1*, which inhibits the Notch pathway (Alcolea et al., 2014), and *Trp53* mutants (Klein et al., 2010; Murai et al., 2018). The distinction between these two possibilities is important since the former involves a neutral scenario where *Pik3ca^H1047R-YFP/wt^* population would exhibit no competitive advantage over wild type but an exacerbated stochastic behavior (i.e. accelerated clone growth but also decline). By contrast, fate imbalance would introduce a selective advantage, cause a net exponential-like growth of mutant clones and lead to mutant cell colonization of the epithelium.

The supralinear growth observed in the average mutant clone size points towards a progenitor fate imbalance in the *Pik3ca^H1047R-YFP/wt^* population (**Figure 3B**). Unfortunately, however, the initial clonal induction efficiency was variable between mice and this precluded reliable confirmation of an overall mutant cell colonization. To circumvent this issue, we decided to explore the distributions of the mutant clone sizes (**Figure 3D**) and extend the maximum likelihood estimation (MLE) approach explained earlier to infer the most likely scenario of mutant progenitor behavior. The following model that allows for a fixed progenitor fate imbalance (Δ) was considered (Klein et al., 2010; Murai et al., 2018):

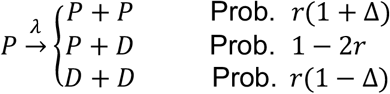

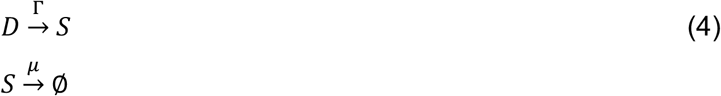

Notice that the neutral-case scenario (balanced fates) corresponds with Δ = 0 (see equivalence to **Eq. 1**), and thus, it can be tested as a nested condition from this model. The assumption on the invariant nature of Δ is made for simplification and becomes reasonable given the low (variable) induction and the relatively short time scales in the experiments; different processes could dampen mutant fate imbalance at longer term (Colom et al., 2020; Murai et al., 2018).

The deterministic ODE equations that describe the time course of the global cell populations from the model in **Eq. 4** are:

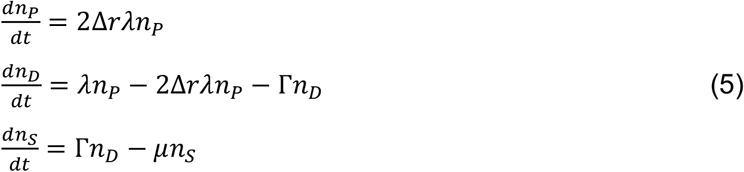

From these, one can infer asymptotical values for and *h* as *t* ≫ *Δrλ*, so that the following relationships between the parameters can be established:

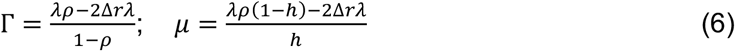

A parameter grid search was done for different values of *r* ∈ [0, 0.5], *ρ* ∈ (0, 1] and Δ ∈ [0, 1] and stochastic model simulations run to estimate theoretical probabilities that clones develop into different sizes *p*_*n*_(*t*, *θ*), as set earlier for the wild type (the mutant average division rate < *λ* > and cell-cycle time *t_cc_* was assumed equal to wild type). By computing the log-likelihood value *l*(*θ* ; *x*) for every set of parameter values θ on the experimental distributions of the number of basal cells per clone we obtained the following parameter estimates (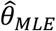 with 95% C.I.) for the *Pik3ca^H1047R-YFP/wt^* progenitors (**Figure 3I**; **Figure S2H**):

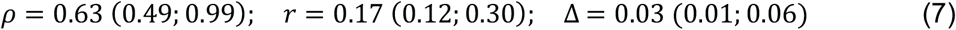

According to the parameter relationships in **Eq. 6**, Г = 4.86 (2.72; 285.48) week^−1^. It follows that independently of the values of *r* and, a neutral SP model with balanced fates (Δ = 0) was significantly less likely to explain mutant progenitor dynamics than a model with fate imbalance (Δ > 0), implying selection (using most favorable parameters for each model, *p* = 0.0002, *** by likelihood ratio test). In fact, the best neutral model found could not produce a good fit on the basal clone size distributions nor explain the curvature in the time course of the average number of basal cells per clone in the mutant population. A SP model with fate imbalance (MLE values from **Eq. 7**) gave an excellent fit on these experimental data (**Figure 3B**; **Figure S2I**). We conclude that *Pik3ca^H1047R-YFP/wt^* progenitor dynamics are characterized by a small, but existent, statistical bias in fate towards an excess of dividing over differentiating daughters per average cell division. This is accompanied by an overall increased proportion of divisions being symmetric, as reflected by the larger value of *r* (**Figure 3I**). Altogether, this would result in individual mutant clones developing into a wider range of possible sizes – more extreme random trajectories – while there is, overall, a relentless colonization of the esophageal epithelium by *Pik3ca^H1047R-YFP/wt^* cell population (**Suppl. Video**).

### 3. Dynamics of suprabasal cells and total clone behavior

A model of *Pik3ca^H1047R-YFP/wt^* selection based on progenitor fate imbalance sets some predictions on the suprabasal component of mutant clones. Therefore, within the limitations of our experimental model (*Pik3ca^H1047R-YFP/wt^* cells could only be detected up to the 1^st^ suprabasal layer, above which the *Pik3ca* locus expression is too close to the background), we turned to study the dynamics of suprabasal and total cell populations as a validation (at least semi-quantitatively) of our model findings.

First, the introduction of a fate imbalance is expected to decrease the relative proportion of suprabasal cells in the mutant clones. In particular, the global suprabasal-to-total cell ratio *h* in the *Pik3ca^H1047R-YFP/wt^* population would decay with the value of *Δr* according to the following rational function (**Figure S2J**) (see **Eq. 6**):

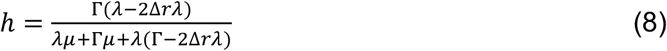

Indeed, the average fraction of 1^st^ suprabasal cells in *Pik3ca^H1047R-YFP/wt^* clones was significantly smaller than in wild type clones across time points (**Figure 3G**). Intriguingly, the experimental data showed a gradual decline in the proportion of suprabasal cells in both genotypes at the late time points, something a simple model with constant parameters would not capture. This apparent departure from homeostasis might be technical but could be related to epithelial changes as mice age (Liu et al., 2019). To achieve a more accurate description of the experimental conditions while retaining the MLE model parameters that suited basal-layer behavior, we thus considered a time-dependent shedding rate μ(*t*), which is the only extra adjustable parameter needed to describe suprabasal cell dynamics. In this way, in the simulations of suprabasal and total clone sizes, μ could be adapted to reproduce the variable value of *h* over time (**Eq. 6**).

Independently of whether we implemented time-adjusted values for μ or used a fixed default value given by the experimental average suprabasal-to-total cell ratio *h*, our (zero-parameter) model fits on the distributions of total (basal + 1^st^ suprabasal) cells per clone were adequate over time points (**Figure 3B**; **Figure S2I**), arguing on the adequacy of the MLE estimates.

Finally, the model predicts differences in the fraction of floating clones (i.e. clones having no basal cells) between wild type and mutant populations at intermediate time points (**Figure S2K**). These differences persisted regardless of whether only P-cells were considered as initially labelled or all populations were initially induced at even proportions. Floating clones reflect transient clones that have lost the proliferative capacity and are to be lost by terminal differentiation. They are expected to be less abundant if progenitors show a fate imbalance favoring proliferating over differentiating progeny. This felt in agreement with trends in the experimental data (**Figure 3H**).

Altogether, mathematical modelling indicates *Pik3ca^H1047R-YFP/wt^* behavior is explained by mutant progenitors showing an increased ability to yield proliferating daughters upon division, which confers this genotype a competitive advantage over wild type cells.

Code used for model simulations in this study can be found at: https://github.com/gp10/DriverClonALTfate

